# Quantification of the Resilience and Vulnerability of HIV-1 Native Glycan Shield at Atomistic Detail

**DOI:** 10.1101/846071

**Authors:** Srirupa Chakraborty, Zachary T. Berndsen, Nicolas W. Hengartner, Bette T. Korber, Andrew B. Ward, S. Gnanakaran

## Abstract

Dense surface glycosylation on the HIV-1 envelope (Env) protein acts as a shield from the adaptive immune system. However, the molecular complexity and flexibility of glycans make experimental studies a challenge. Here we have integrated high-throughput atomistic modeling of fully glycosylated HIV-1 Env with graph theory to capture immunologically important features of the shield topology. This is the first complete all-atom model of HIV-1 Env SOSIP glycan shield that includes both oligomannose and complex glycans, providing results which are physiologically more relevant than the previous models with uniform glycosylation. This integrated approach including quantitative comparison with cryo-electron microscopy data provides hitherto unexplored details of the native shield architecture and its difference from the high-mannose glycoform. We have also derived a measure to quantify the shielding effect over the antigenic protein surface that defines regions of relative vulnerability and resilience of the shield and can be harnessed for rational immunogen design.

## Introduction

Protein glycosylation is an essential aspect of post-translational modification, with 50-70% of human proteins been estimated to be glycosylated to some degree(An et al., 2009). These glycans play significant roles in numerous biological processes, such as cellular signaling, recognition and adhesion, protein folding, structural stability, and immune system interactions(Dennis et al., 1999; Dwek, 1996; Moremen et al., 2012). Additionally, envelope proteins from several high-risk viral pathogens which hijack the host protein production and glycosylation machineries, such as HIV (lentivirus)(Burton and Mascola, 2015), Coronavirus (Vankadari and Wilce, 2020; Walls et al., 2016), Lassa (arenavirus)(Sommerstein et al., 2015), Hepatitis C (flavivirus)(Zhang et al., 2017), Epstein Barr (herpesvirus)(Szakonyi et al., 2006), Ebola (filovirus)(Ilinykh et al., 2018; Lennemann et al., 2014), and Influenza(Wang et al., 2009) are also heavily glycosylated. These surface-expressed viral proteins are important immunological targets for neutralizing antibodies that can block viral infection of cells, and form the primary focus of vaccine studies(Amanat et al., 2018; Saphire et al., 2018). However, the dynamic and dense glycan coating can effectively act as a shield for the underlying envelope protein, masking antigenic surfaces, barricading it from the adaptive immune system, and defending against immune surveillance(Crispin et al., 2018; Doores, 2015; Karsten and Alter, 2017; Ringe et al., 2019; Stewart-Jones et al., 2016; Wagh et al., 2018). A deeper molecular level understanding of the glycan shields in these pathogenic viruses may help inform vaccine design strategies that can overcome this protective barrier.

The HIV-1 Envelope Glycoprotein (Env) is a heterodimeric trimer composed of proteins gp120 and gp41, and is responsible for the molecular recognition of the host receptor and fusion into host target cells. A number of obstacles hinder traditional vaccine design methods in case of HIV-1 – namely its remarkable sequence diversity, conformational plasticity, dramatic shifts in position and number of glycans(Zhang et al., 2004), and extremely dense glycosylation making up to approximately half the mass of the entire Env molecule(Lasky et al., 1986). As a result, there has been only limited success in eliciting broadly neutralizing antibodies (bNAbs) to Env vaccine immunogens(Bradley et al., 2016; Haynes et al., 2014; Walker et al., 2010). Structures of bNAbs in complex with Env indicate that these antibodies need to extend through the glycan shield in order to engage with the epitopes at the protein surface(Huang et al., 2014; Julien et al., 2013; Lyumkis et al., 2013; Pancera et al., 2014). Some bNAbs have evolved to include conserved glycans as part of the epitope(Crispin and Doores, 2015; Falkowska et al., 2014; Mouquet et al., 2012; Walker et al., 2009). Moreover, these surface glycans are also critical for Env folding, viral assembly, and infectivity.

Almost all the glycans on the Env are N-linked and occur at potential N-glycosylation sites (PNGS, given by the sequon Asn-X-[Ser or Thr], where X is not proline)(Stanley et al., 2015). Such glycans usually have about 10-20 pyranose rings covalently connected in a tree-like structure to the asparagine(N) residue of the sequon. These can further undergo additional processing(Crispin and Doores, 2015) into complex sugars from simple oligomannose, depending on their spatial location, local protein content, crowding, and producer cell type. A high degree of processing is indicative of exposure or accessibility of sugars to enzymes. At regions with dense crowding of glycans, steric constraints limit the activities of the carbohydrate processing enzymes(Pritchard et al., 2015a; Pritchard et al., 2015b) as the proteins fold and translocate through endoplasmic reticulum and Golgi apparatus.

The extreme dynamics and conformational heterogeneity stemming from large variance of possible constituent sugars, linkage, branching patterns, and rotamer flexibility, make the study of these glycans immensely challenging(Imberty and Perez, 2000; Walsh, 2010). Traditional X-ray crystallography methods are rendered ineffectual(Chang et al., 2007) since this heterogeneity prevents proper crystallization of the glycoproteins. Inherent glycan flexibility also complicates analysis by ensemble techniques like Nuclear Magnetic Resonance (NMR) and single-particle electron cryomicroscopy (cryo-EM) where the signal from glycans is often too noisy for structural interpretation at the resolutions necessary for building atomic structure. (Chang et al., 2007; Davis and Crispin, 2010; Slynko et al., 2009; Woods et al., 1998). Within the ~300 experimental structures of HIV-1 envelope glycoprotein in the Protein Databank (PDB) (Berman et al., 2000), only a quarter of the total glycan content has been structurally resolved and are mostly in high mannose form. Recently there have been a few cryo-EM and X-ray structures published of natively glycosylated Env(Barnes et al., 2018; Gristick et al., 2016; Lee et al., 2016). Since most of these structurally resolved glycans were stabilized by interaction with antibodies, they provide little information about the structural and dynamic details of the individual and collective glycans. Furthermore, all computational studies of HIV-1 Env glycosylation comprise of oligomannose glycan moieties ranging from mannose-5 to mannose-9(Ferreira et al., 2018; Lemmin et al., 2017; Liang et al., 2016; Stewart-Jones et al., 2016; Yang et al., 2017). The main drawback affecting the quality of such MD simulations is the robustness of conformational sampling. Due to the intrinsically dynamic nature of glycans, to effectively sample a biologically relevant energy landscape of the glycoprotein, long and often multiple trajectories need to be run, preferably with enhanced sampling techniques(Yang et al., 2017).

We had previously established a high throughput pipeline to robustly build atomistic models of glycans, sample the glycan conformational space, construct the glycan network topology, and extract molecular level description of the viral glycan shield (Berndsen et al., 2019). We had also set up a protocol to experimentally validate the models by generating synthetic cryo-EM datasets and comparing the resulting maps in a quantitative manner. Here, we have employed a similar quantitative comparison using computational and experimental single-particle cryo-EM data and show that the dynamics of high-mannose and native glycosylation remains practically indistinguishable around the glycan stem β-mannose (BMA) residue. This provides further rationale towards the need for computational modeling of complete glycans in a natively glycosylated shield. Presently this seems to be the only way to elucidate the ramifications of such complex sugars on the shield architecture. Therefore, we have built a glycosylated Env having both oligomannose and complex glycans(Behrens et al., 2016; Cao et al., 2017) based on the information provided by mass spectrometry (MS)(Behrens et al., 2016). To our knowledge, this is the first time that a complete computational model of Env SOSIP was generated to include processed glycoforms, thus obtaining results which are immunologically more relevant.

Using these atomistic models, we have employed graph theory to capture the glycan shield topological network, pinpoint potential interaction pathways, and identify concerted behavior of the glycans. The glycan-glycan and glycan-protein interactions influence the behavior of the shield as a whole and can affect distant sites in the glycan network(Behrens et al., 2016; Ferreira et al., 2018; Stewart-Jones et al., 2016). Analyses of various network attributes, such as relative centrality of different glycans and critical subnetwork properties, have aided in detailed examination of the native glycan shield in the context of bNAb interactions. Important global and local feature differences due to native-like glycosylation pattern come to light as compared to all high-oligomannose glycans, arising from the presence of charged sialic acids at the tips and fucose at the base. Both the simulated models and cryo-EM maps identify a change in projection angle relative to the peptide surface for fucosylated glycans. We have also been successful in quantifying the glycan shield based on the number of sugar heavy atoms encountered over the antigen surface that can be implemented to define regions of relative vulnerability and regions where the glycan shield blocks access. Due to the rapid yield rate of this method, it can be applied to a large number of diverse HIV-1 sequences or can be seamlessly extended to model glycan shields of other viruses in a rapid manner.

## Results

### Selection of site-specific glycans for native glycosylation of HIV-1 Env

The soluble, recombinant BG505 SOSIP.664 (here on referred simply as BG505) trimer has been well characterized as a native-like, Env-mimetic model, and serves as the prototypical immunogen in several vaccine development programs(Sanders et al., 2013; Sanders et al., 2015). For our study, we have used this system in the pre-fusion closed conformation, for which several structures have been determined(Barnes et al., 2018; Gristick et al., 2016; Lee et al., 2016; Stewart-Jones et al., 2016). A single protomer of this BG505 protein contains 28 N-linked glycosylation sites. While all N-glycans share a common core sugar sequence, they are broadly classified into three types (**Figure 1**): (i) oligomannose, in which only mannose residues are attached to the core; (ii) complex, in which differently branching “antennae” are attached to the core; and (iii) hybrid, which is a mix of the first two types. Recent novel MS-based approaches have been successful in identifying site-specific glycosylation profile in multiple HIV-1 trimers(Cao et al., 2017; Go et al., 2017). It is known that each glycosylation site at BG505 has a distribution of multiple possible glycan species, depending on various factors, such as type of cell lines, purification methods and sources, structural constraints, and so on. We used a site-specific distribution of glycan type at each PNGS that was obtained by Behrens *et al*(Behrens et al., 2016) and Cao *et. al.*(Cao et al., 2017), to identify the most likely glycan species at each site. The glycan species at each site was selected based on the highest relative abundance per the MS studies (Figure 1 in reference (Behrens et al., 2016)), and is listed in Supplementary Information (SI) **Table S1**. The structure schematic for each of the selected glycan species have been shown in **Figure 1A**.

**Figure 1:**
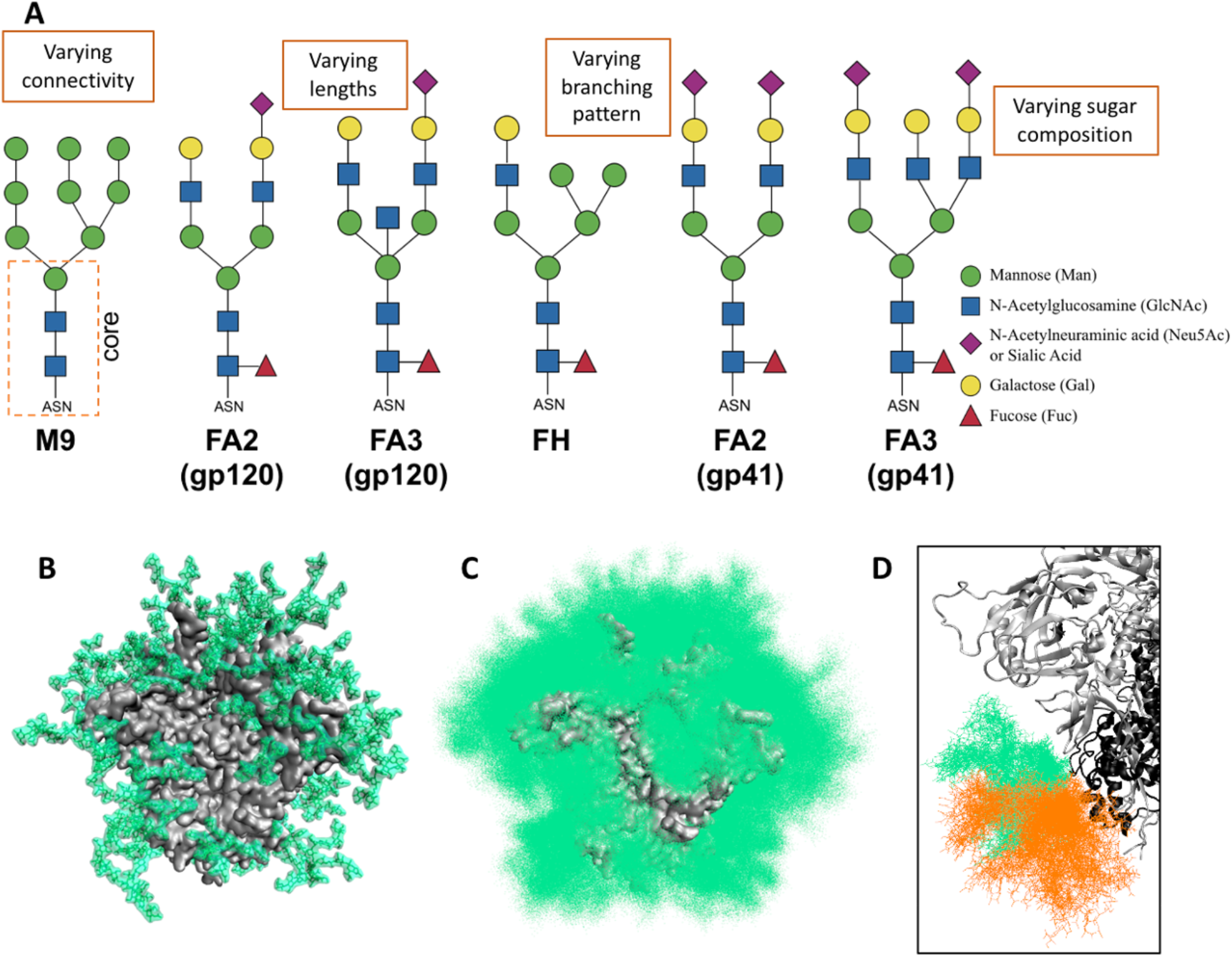
Native glycosylation and shielding of HIV-1 Env. (A). Schematic representation of N-glycan types. The common core of Man3GlcNAc2Asn is indicated. The glycan species selected for different sites of the BG505 native model (as given in **Table S1**). M9 ≡ mannose-9, FA2 ≡ fucosylated two-antennae, FA3 ≡ fucosylated three-antennae, FH ≡ fucosylated hybrid. Complex glycans in gp41 are different from those in gp120, as per site-specific mass spectroscopy experiments. (B) Single snapshot of modeled natively glycosylated BG505 structure, with each glycan taking up a particular conformation. (C) Cumulative shielding effect over time due to the dynamic nature of the glycans. 100 randomly selected models from the 1000-structure ensemble is shown here. (D) Neighboring glycans within close proximity can sample overlapping volumes. Glycan ensembles N88 shown in green and N625 shown in orange.

### High-throughput atomistic modeling of the native glycan shield

In the static pictures as obtained from fully glycosylated structures(Gristick et al., 2016; Stewart-Jones et al., 2016) and even single snapshots from computational models, a significant fraction of the protein surface appears to be exposed, with each glycan taking up a particular conformation (**Figure 1B**). However, due to their flexible and dynamic nature, these glycans are not confined to a particular conformation, instead sampling a large volume in space. A previous simulation study suggested(Guvench et al., 2011) that the root mean squared deviations (RMSD) for the carbohydrate regions are more than 4 times larger than that of the underlying protein loops. As a result of this conformational variability, the cumulative effect of the glycans over time is like that of a cloud of glycan atoms that shield the underlying surface from any approaching protein probe (an antibody for example) as illustrated in **Figure 1C**.

Here we have incorporated experimental information-driven conformational sampling in the flexible unstructured protein regions to model the entire glycan shield. Details of the modeling procedure are given in the Methods section, and are briefly reiterated here. A robust ensemble of BG505 glycoprotein atomic models was generated by utilizing a template-free glycan modeling pipeline including a sequence of refinement steps with restraints to enforce proper stereochemistry. The underlying protein scaffold was built by utilizing templates from a number of experimentally determined Env structures deposited in the Protein Data Bank by assimilating the differences in local structural regions including loops, that arise due to flexibility. The glycans were modeled *ab initio* on several distinct protein scaffolds by implementing the ALLOSMOD(Guttman et al., 2013; Weinkam et al., 2012) package of MODELLER(Eswar et al., 2006; Sali, 1995) in a streamlined pipeline.

Due to the initial randomization of the glycan orientations, this integrated technique can sufficiently sample a physiologically relevant conformational space accessible to carbohydrates in a very short time, leading to the overall spatial shielding effect (**Figure 1C**). The torsion angle distributions of 10 different inter-glycan linkages within the ensemble was compared with those obtained from different glycan structures available in the PDB database using GlyTorsion webserver(Lütteke et al., 2005) and is shown in **SI Figure S1**. These distributions match very well between our generated ensemble and the PDB structures (see SI for details), indicating that the modeled glycans sample a stereo-chemically relevant landscape. In order to understand the effects just due to native glycans, we have also built a similar ensemble with mannose-9 glycans at all glycosylation sites, (hereon referred as all-man9 model) for comparison.

### Quantitative comparison of simulated models with cryo-EM data indicates glycoform-independent glycan stem dynamics

We generated simulated cryo-EM density maps for both the native and all-man9 ensembles by utilizing the method that we established earlier to convert *in silico* atomistic models to synthetic cryo-EM data(Berndsen et al., 2019). Then, we quantitatively compared them to the experimental cryo-EM maps of BG505.SOSIP.664 from HEK 293F cells which produce a native-like glycan shield including both oligomannose and complex glycans, and from HEK 293S cells which do not produce complex glycans(Berndsen et al., 2019). The experimental and simulated maps show similar features in the shield, with only the first few sugar residues at each site defined at high resolution (**Figure 2A**). A scale-space analysis developed in our earlier study(Berndsen et al., 2019) also shows a similar trend to the experimental data (**Figure 2B**). Progressive smoothing of the map is accompanied by a large expansion in glycan volume that plateaus around 1.75-2 SD. The effect of the glycans is best appreciated in comparison to a map generated from a non-glycosylated ensemble (**Figure 2B - dashed line**). This suggests the simulation is capturing native-like dynamics in the glycan shield, at least globally. The 1.75 SD Gaussian filtered maps are visualized at high and low thresholds in **Figure 2C**, revealing a similar overall shape and topology. At the level of individual glycan dynamics, comparison of the normalized mean intensity at the location of each BMA residue for the simulated map with that of the experimental BG505_293F map reveals a strong positive correlation (**Figure 2D-E**), with a correlation coefficient of ~0.82 (p=6.82e-6). We also confirmed that local map intensity around BMA residues accurately captures relative differences in Root Mean Squared Fluctuations (RMSF) at each glycan (**SI Figure S2**) for the native ensemble.

**Figure 2:**
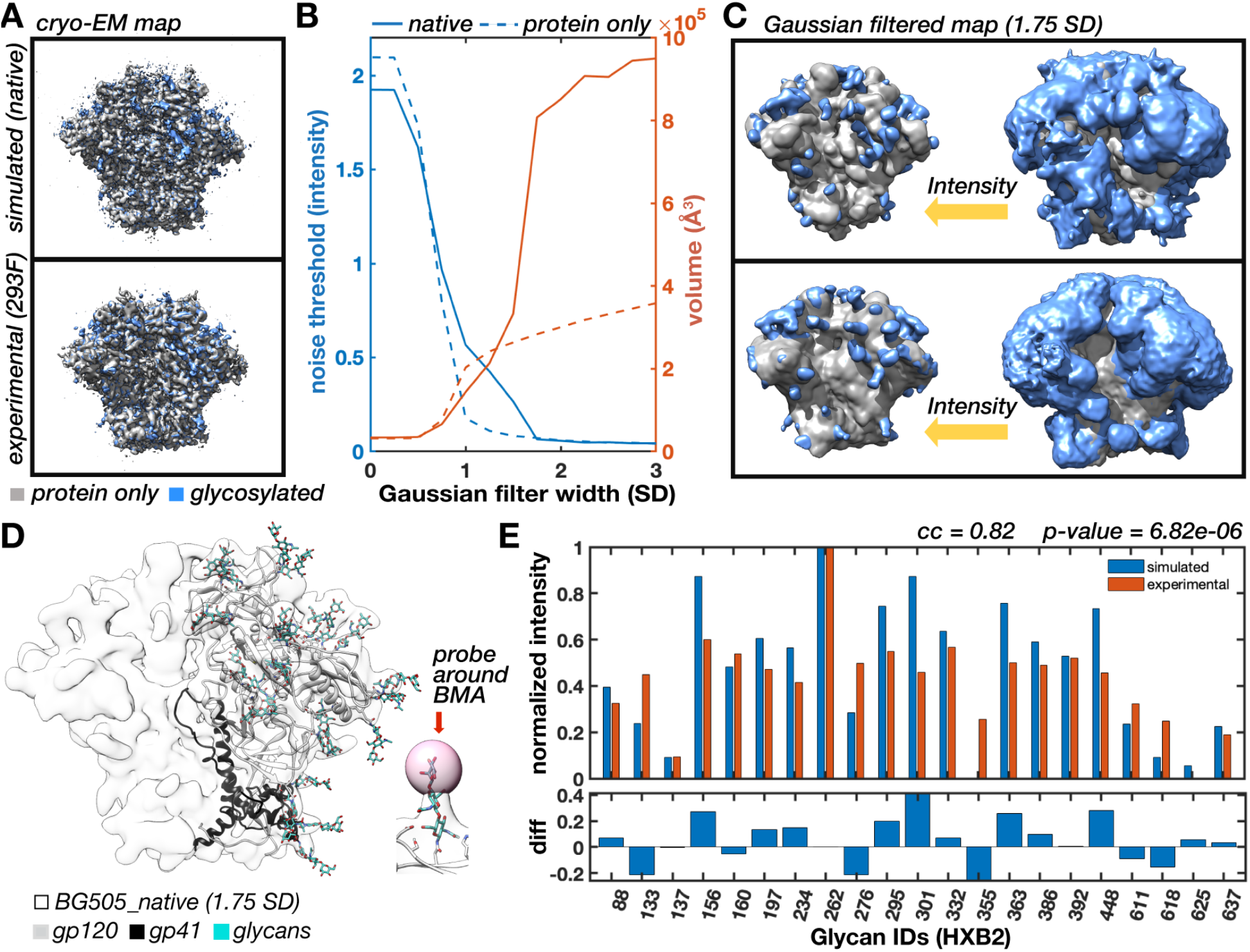
Comparison of glycan dynamics between experimental and simulated cryo-EM maps. (A) Experimental BG505_293F (top) and simulated BG505_native (bottom) cryo-EM maps sharpened and filtered at the global resolution (~3Å), with maps of de-glycosylated trimers displayed underneath to highlight the glycan density. The BG505_293F data is reproduced from (Berndsen et al., 2019) and density corresponding to the three RM20A3 Fabs bound to the base of the SOSIP were deleted for the sake of aesthetics (B) Noise threshold and map volume at the noise threshold as a function of Gaussian filter width (standard deviation - SD) of the BG505_native and de-glycosylated maps showing the progressive emergence of glycan signal at lower resolution. (C) 1.75 SD Gaussian filtered maps from panel A displayed at high and low isosurface thresholds. (D) Gaussian filtered BG505_native map with relaxed atomic model used for measuring local glycan dynamics along with a close up of a single glycan with a spherical probe around the BMA residue. (**E**) Bar plot showing relationship between the normalized site-specific mean local intensity from the BG505_293F (experimental) and BG505_native (simulated) maps along with the absolute difference at each site along with the Pearson correlation coefficient.

An interesting outcome of the above comparison is that the correlation coefficients remain similarly strong not only when we compare the all-man9 model to BG505_293S data, but also when the native model is compared to high-oligomannose experiments and vice versa (**SI Figure S2**). Therefore, while these comparisons give credence to the validity of the model in accurately capturing the relative position and flexibility of the glycan stems, they also indicate that the glycan dynamics are practically identical within the observational limits of cryo-EM analysis between complex and oligomannose glycans. The native glycosylation pattern on the shield topology only manifest at the level of the flexible antennae beyond the stem and can therefore only be observed in detail using computational models.

### Fucose-mediated changes in glycan orientation distinguish native glycan shield

While the glycan stem shows indistinguishable dynamics at the BMA sugar, an important difference between complete models of complex and all-man9 arise due to the presence of the fucose ring at the core of the complex sugars (**Figure 3**). By structurally aligning the first core sugar and the three residues at the base of the glycan (n-1, n, n+1; where n is the glycosylated asparagine residue) for all the structures in the ensemble, the oligomannose glycans are found to be spaced symmetrically around the core, while complex sugars have a distinct bend away from the side where the fucose ring is present (**Figure 3A**). The change in projection angle of the principal axis of the glycan relative to the protein surface is predicted to be ~10° in our models. This difference in orientation preference, along with the presence of the negatively charged sialic acid tips of the antennae, play notable roles that govern the differences between high oligomannose and native glycosylation.

**Figure 3:**
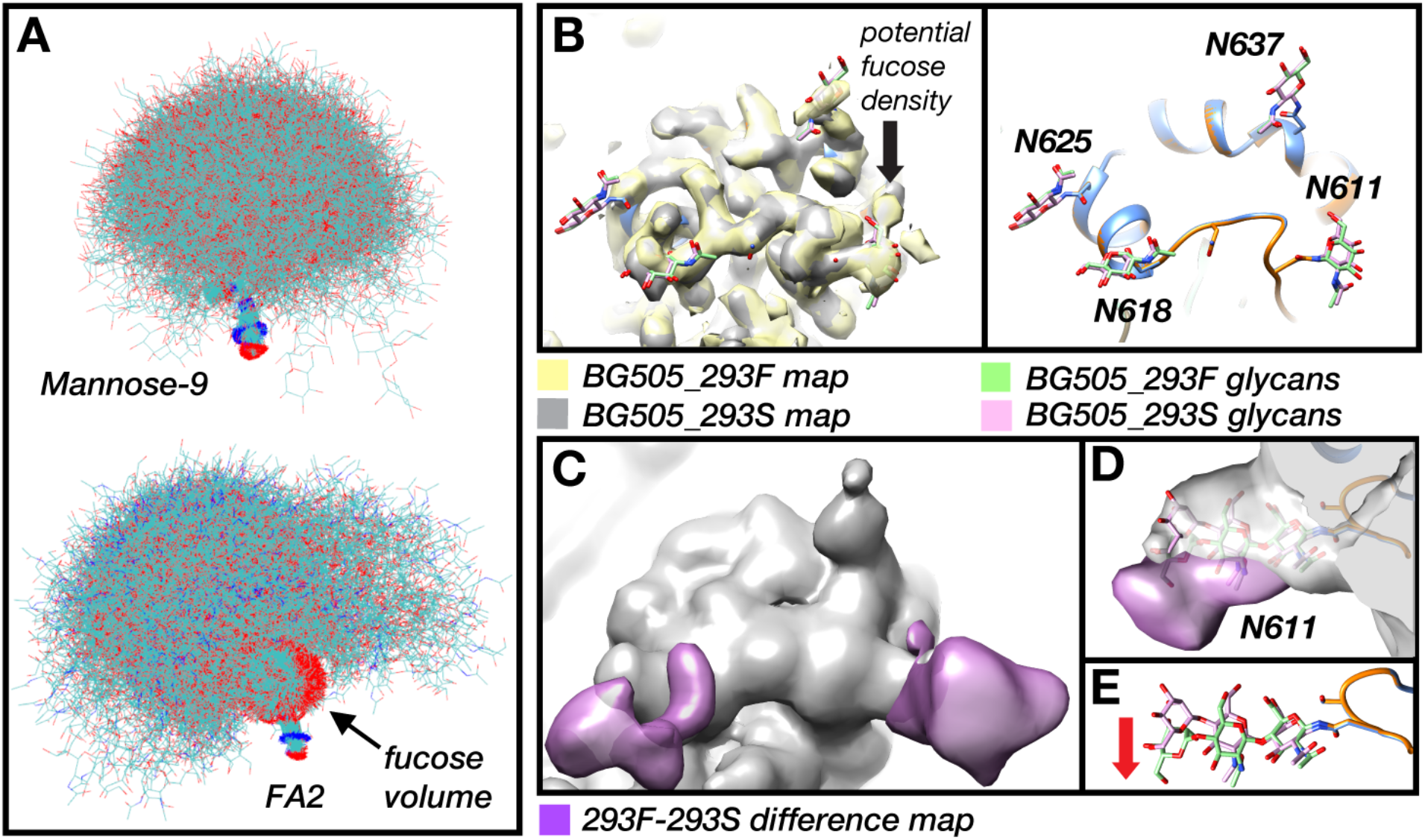
Model and Experimental cryo-EM data reveals subtle change in orientation due to core fucose. (A) Difference in overall glycan orientation between all-man9 and fucosylated complex glycan in simulated model. The first core sugar and three residues at the base of the glycan (n-1, n, n+1; where n is the glycosylated asparagine residue) were aligned for a single sugar ensemble. Oligomannose samples a space symmetrically around the core, while, complex sugars have a distinct bend away from the side where the fucose ring is present. (B) Cryo-EM maps and refined atomic models for BG505_293F and BG505_293S (add citation) showing the 4 glycans from gp41. Only the core NAG residues can be confidently built into the sharpened maps. (C) Gaussian filtered BG505_293S map (gray) and BG505_293F-BG505_293S difference map (purple). (D) Close up of the glycan at N611 showing asymmetric difference signal around the glycan stem. (E) Full glycan stalks built and relaxed into both Gaussian filtered maps showing more pronounced change in projection angle.

The change in projection angle from the fucose ring manifests as a reasonably large change at the tips of the antennae, but the overall change in position at the core N-acetylglycosamine (NAG) is too small to be captured in the high-resolution cryo-EM maps (**Figure 3B**). However, a close examination of the low-resolution difference map between the BG505_293F and BG505_293S maps (Berndsen et al., 2019) does reveal some evidence for such changes in projection angle at sites with complex glycans (**Figure 3C-E**). For instance, the difference signal around the glycans at N611 and N618 is asymmetric around the glycan stem and is therefore likely not the result of differences in occupancy. The difference signal at N611 is located primarily on the underside of the glycan stem, on the face opposite where we identified density likely belonging to the fucose ring (**Figure 3D**), which is suggestive of a change in projection angle similar to what we observed in the simulations. In addition, examination of the glycan stem models built and relaxed into the Gaussian filtered BG505_293F and BG505_293S maps used for assessing local map intensity (**Figure 3E**) show a slight change in projection angle, further confirming this fucose-mediated glycan reorientation.

### Network analysis of glycan topology explicates the shield connectivity

Network analyses has historically been used to study protein allosteric frameworks and evolutionary paths (Beleva Guthrie et al., 2018; Eargle and Luthey-Schulten, 2012; Huang et al., 2017; Sethi et al., 2013; Skjaerven et al., 2014), and only recently begun to be applied to in glycoprotein structural characterization (Lemmin et al., 2017). While each glycan exerts effects in its immediate vicinity, due to the inter-glycan interactions, their influence can percolate over long distances across the surface of protein(Berndsen et al., 2019). Long-range glycan interactions can occur, as with perturbation of a glycan at one site can affect the processing and antibody interactions of another glycan at a distant site(Behrens et al., 2016; Bricault et al., 2019; Doores, 2015; Stewart-Jones et al., 2016). We have previously shown that understanding long-range interactions between glycans is important for characterizing global properties of the glycan shield(Berndsen et al., 2019).

Now, we implement the graph-theory based approach to native glycan shield. Details of modeling the network are recapitulated in the Methods section. The natively glycosylated BG505 trimer, like all HIV-1 Envs, has a highly dense glycosylation pattern and each glycan samples a particular region in space (**SI Figure S3A-B**), such that neighboring glycans can occupy overlapping regions (**Figure 1D**). The fraction of volume overlap gives a measure of the interaction probability between the constituent glycans (**SI Figure S3C**). Within a protomer there are three main regions of overlap – the apex, the gp41 base, and the glycan-dense central high-mannose patch (HMP). Inter-protomer overlaps are primarily due to V1 and V2 loop glycans near the trimer apex. **Figure 4A** shows the obtained network of BG505 native glycosylation unfolded and laid out in two dimensions for the ease of visualization. This is a force-directed layout, which uses attractive forces between adjacent nodes and repulsive forces between distant nodes to reach an optimum distribution of the node points in space(Fruchterman and Reingold, 1991). For conceptualization, the network relative to the BG505 structure is provided in **SI Figure S3D**.

**Figure 4:**
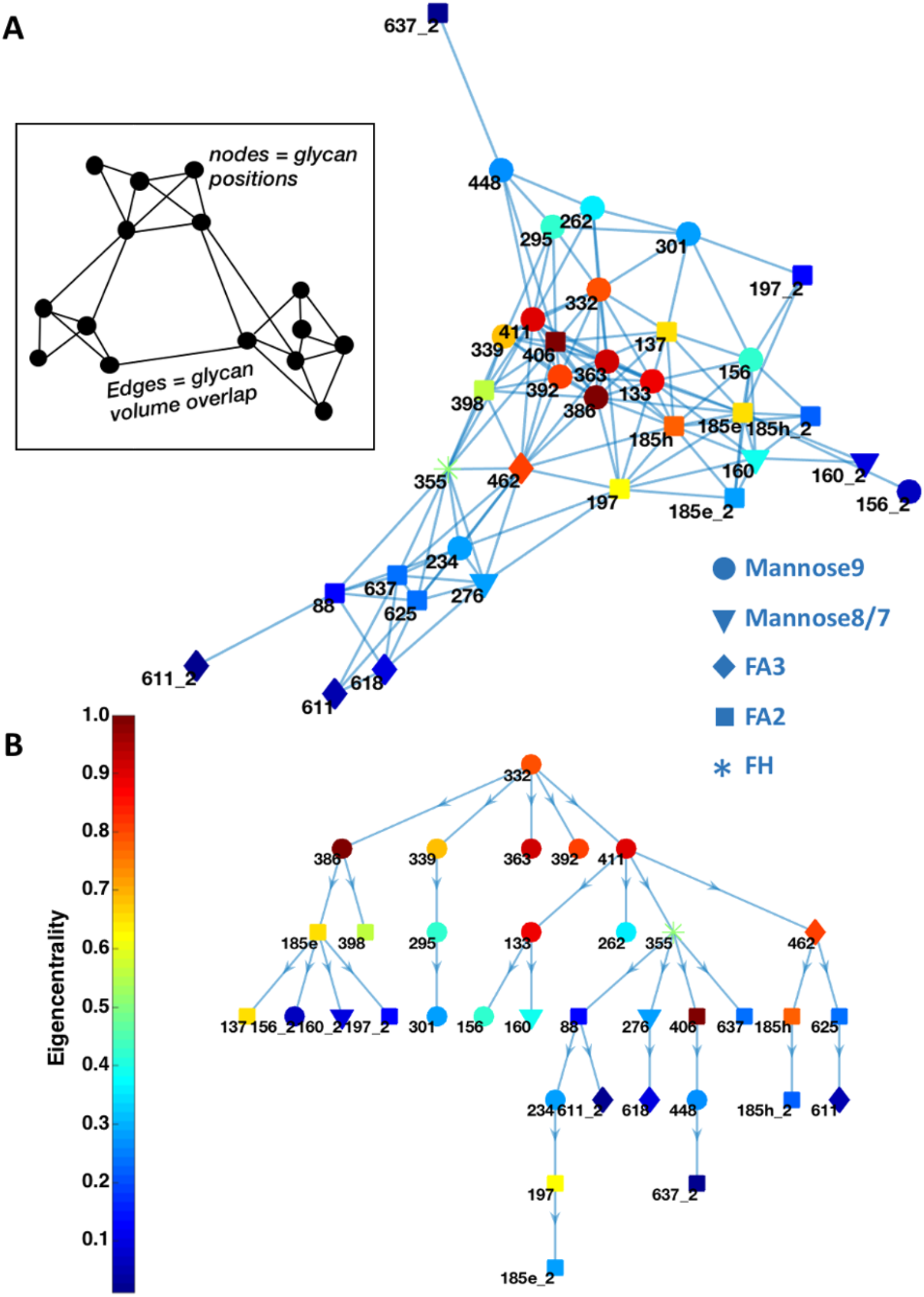
Spatial network of BG505 native glycosylation. (A) Visual representation of network in 2-dimensions, based on a force-directed layout. Each glycan forms a node-point on the graph, two nodes are connected by an edge (inset, from reference (Berndsen et al., 2019), Figure 8). The nodes are colored by the normalized eigenvector centrality of the glycans. (B) Shortest path of communication between glycan N332 and all other glycans in the network.

The nodes in the central region around the V4 loop, in the HMP, are very highly inter-connected. Glycans in the V1, V2 apex are also reasonably well-connected, but the connections at the base near gp41 are relatively sparse, both within the locality, and globally with the rest of the network. Glycans at positions 234, 355, 462 and 276 connect the sparse base region of the network with the dense apex and central regions. Glycan 156, 160, and those in the V2 loop, enable inter-protomer glycan interactions at the apex, stabilizing the protein quaternary trimer structure(Berndsen et al., 2019). Glycan 197, crucial in the binding of CD4bs and V3-specific antibodies(Li et al., 2008; Liang et al., 2016), connects the central crowded region of the network between neighboring protomers. gp41 glycans 611 and 637 also contribute to inter-protomer pathways, with glycan 637 communicating with glycan 448 directly in the central mannose patch.

### Utilizing the glycosylation network to extract functionally relevant topological features

The native glycosylation network is well-connected as indicated by the network degree, or the number of connections a node has to other nodes (**SI Figure S3E**). Due to the high structural fluctuations of the V2 loop, the glycans on and flanking V2 can interact with a number of other neighboring nodes, having the highest degrees of connectivity. There is dense crowding of glycans around the high mannose patch with a number of inter-connections, resulting in the glycans of this patch being less processed because of reduced accessibility to enzymes(Coss et al., 2016; Doores et al., 2010; Pritchard et al., 2015a). The network degree values are relatively lower around the base of the protein on and around gp41, reflective of the sparse architecture of glycan topology in this region. The site N332 is central in the glycan interaction network, and is known to have both direct and subtle long-range influences over a number of bNAb sites(Behrens et al., 2016; Bricault et al., 2019). We calculated the shortest paths to all glycans from the N332 glycan site using the Floyd-Warshall algorithm(Floyd, 1962; Warshall, 1962), where the glycan distances were edge-weighted (**Figure 4B**). Our calculation shows the most probable pathway over which each glycan feels the influence of the glycan at site N332, providing useful insights into how these inter-glycan communications can occur.

To elucidate the relative influence of each of the glycans on the network, we calculated the eigenvector centrality, or eigencentrality, of the nodes to measure its connectivity to the network. A large relative value indicates that the node is well connected to the network and thus is “centrally” located, whereas a low relative value indicates that the node is on the periphery of the network. Operationally, the eigencentrality is the eigenvector associated with the largest eigenvalue of the adjacency matrix, that in position (i,j) reports the overlap of the ensemble of glycan at sites i and j. Value of that particular matrix element will be zero if the glycans in position i and j do not physically interact (details given in Methods section). The normalized eigencentrality of the glycans are projected on the network as a colormap in **Figure 4A**. The eigencentrality increases towards the middle of the graph, with the glycans at the crowded HMP with a large number of connections between each other having the highest centrality values. When we increase the threshold of volume overlap needed to form an edge, those glycans with the least centrality leave the network first. Glycan 611 is the first to be eliminated, succeeded by some of the inter-protomer interactions and the other gp41 glycans, following the relative centrality scores. The HMP with high eigencentrality values persists throughout even with high overlap thresholds. We have previously confirmed for all-man9(Berndsen et al., 2019) that successive enzymatic digestion of glycans from Env loosely follows a pattern that matches with the network centrality. Those glycans which are sparsely connected to each other, having lower centrality, are eliminated earlier during the process of endoH digestion. On the other hand, the glycans having higher network centrality, such as those in the HMP, takes longer to be eliminated by the digestive enzymes. Thus, the eigenvector centrality provides a measure of the crowding of the highly central glycans which makes them difficult for endoglycosidase to access.

### Influence of complex glycans on overall network topology

The degree of connectivity decreases almost throughout the network for the native glycosylation (**SI Figure S4A**) compared to all-man9. The number of stable glycan-glycan interactions at several glycan sites decreases because spatially proximal glycans avoid unfavorable charge-charge interactions. For example, the uncharged glycans 234 and 355 takes up a position central to all the neighboring charged sugars, increasing the distances between them, and the charged N406 glycan buries itself in the middle of the surrounding high-mannose glycans (**Figure 4A**), clustering together with them. In fact, removal of this charged glycan N406 glycan can increase the processing of the neighboring high-mannose moieties(Cao et al., 2018). Further evidence for such alterations in glycan connectivity due to the presence of complex glycans can be observed in the comparisons of the RMSF variations. A detailed analysis has been presented in the SI and illustrated in **SI Figure S5**. Combination of the network node location differences and RMSF variations suggest that the presence of the charged sialic acid at the tips of these complex glycans can dictate the interactions between neighbors, increasing their structural variations if surrounded by other charged glycans, or reducing them if a stable conformation buried between uncharged high-mannose patches can be found. This overall decrease in connectivity slightly increases the diameter of the native network, increasing the mean number of hops for the shortest path from 2.4 hops (and distance 0.19) in the all-man9 model to 3 hops (and distance 0.27).

The distribution of centrality also shifts between the two glycosylation models (**SI Figure S4B**). The centralities of the glycans at 137, 234, 355, 398, 406, 462 and those at gp41 increase due to native glycosylation. On the other hand, the V1 and V2 loop glycans as well as 262, 295, 363 decrease from the all-man9 model. **Figure 5A** illuminates the difference in adjacency matrices between the two models, with blue color indicating decrease in edge weight, and red indicating increase in edge weight in native network, as compared to all-man9. Overall connectivity goes down in the apex and the central patch for the native, and increases within gp41 glycans, as compared to all-man9. The differences between the two networks (with at least 10% edge-weight change) are shown in **Figure 5B** and **C**. The pathways around 197, as well as the inter-protomer interactions involving 156, 185e and 637 become stronger in native glycosylation. (**Figure 5B**). Some of these 197 connections, such as those with 276 and the V5 loop glycans occur across the CD4 binding site. It was previously shown that in predominantly high-mannose Env structures, CD4bs targeting antibody VRC01 has very little interaction with glycans 197 and 276(Stewart-Jones et al., 2016). Conversely, in another Env structure with fully processed native glycans, a VRC01-like antibody called IOMA interacted extensively with both 197 and 276(Gristick et al., 2016). This matches our observation of increased orientation of glycan 197 over the CD4bs in native glycosylation pattern. However, the paths connecting 137, 156, 301, and 197 (of neighboring protomer) are significantly weakened in the native model (**Figure 5C**). Subnetworks involving V4 loop glycans and some of the central mannose patch become weaker due to the presence of native glycans.

**Figure 5:**
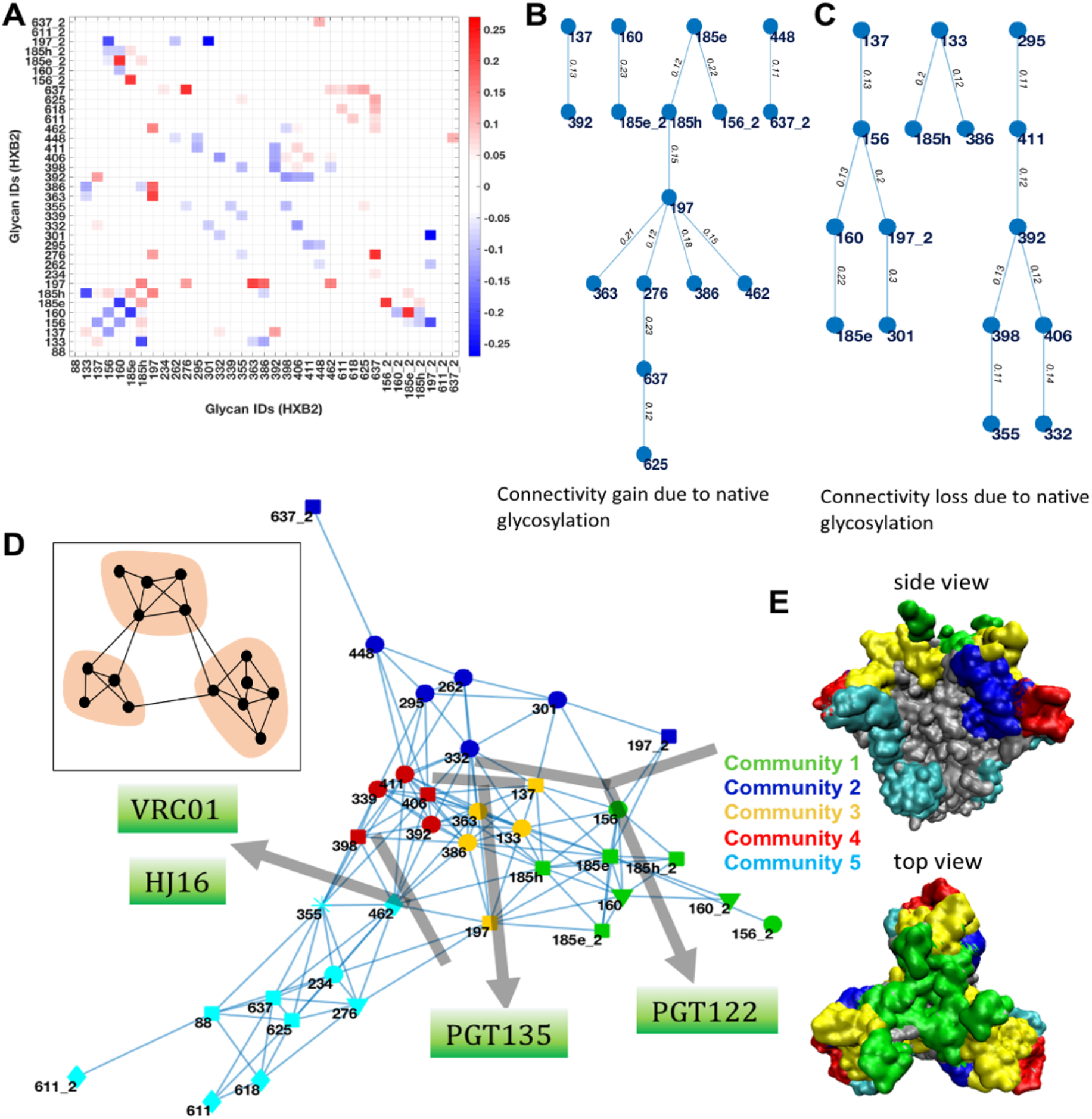
Network difference and community analysis. (A) Difference in adjacency matrices between native and all-man9 models (native minus all-man9). Blue color indicates decrease in edge weight, and red indicates increase in edge weight in native network, as compared to all-man9. (B) Increase and (C) decrease in connectivity due to native glycosylation in comparison with all-man9 model (only those edges that change by 10% or more are shown). (D) 5 different communities were identified based on modularity maximization. The sub-community junctions identify susceptible regions in the shield where the shown HIV antibodies tend to bind. (E) Location of each community projected on the Env surface.

### Community analysis to determine breaches in glycan shield

In order to interact and neutralize the Env protein, antibodies look for breaches in the shield where glycan densities are sparse. This information of relative sparsity is encoded within the glycan network, in terms of edge clustering. We examine this clustering of glycan interactions within the network by defining communities using a modularity maximization approach that divides the network into sub-modules or groups (see Methods). Communities have high density of edges between the nodes within a module and comparatively sparse edges connecting different modules (**Figure 5D inset**). This analysis identifies five distinct communities within the BG505 native glycan network (**Figure 5D and E**). The apex glycans around loop V2 from the three protomers together form a single community (1, green). Right below that, glycans 262, 295, 301, 332 and 448 (2, blue) forms a community around the V3 and alpha 2 helix. Glycans 133, 137, 197, 363, and 386 (3, yellow) and 339, 392, 398, 406 and 411 (4, red) form two distinct communities involving the glycans on and surrounding V4 loop region. Rest of the glycans 88, 234, 276, 355 and 462 from gp120 and all four glycans from gp41 form the fifth community (5, cyan), though the modularity value is lower due to sparser connections. The possibility of an approaching probe reaching the protein surface through these strongly connected communities is low. A similar study clustering glycans from microsecond simulation runs of BG505 SOSIP was performed by Lemmin *et. al.*(Lemmin et al., 2017), identifying 4 glycan microdomains that roughly correspond to our modules 1, 2, (3+4) and 5. However, in that study, mannose-5 glycoform was used at all sites, and due to the smaller length of these glycans, some of the interactions, including inter-protomer interactions were not observed. Remarkably, in that study, junctions between microdomains were found to indicate regions of relative vulnerability. The communities we identified, also demarcate the regions where the glycan shield can be penetrated. Broadly neutralizing antibodies whose binding epitopes are known target these community boundaries (**Figure 5D**). Thus, glycan community dynamics can help to determine susceptible regions of the glycan shield, and can be further used for guiding immunological studies.

### Quantifying the vulnerability of glycan shield for antibody-based neutralization

To capture the impact of the glycan shield acting as an immunological barrier, we quantified the sugar barrier over the Env protein surface using the ensemble of structures that have been generated. Towards this goal, we have defined the glycan encounter factor (GEF) as the number of glycan heavy atoms encountered by an external probe approaching the protein surface. This parameter is calculated at each residue on the surface of the protein within a probe of diameter 6Å calculated using our ensemble. The highest value obtained is 14 for the native model, and is located in the HMP. Based on existing structures, the main interaction points between Env and bNAbs are often hairpin-like loop regions. Even large-scale atomistic simulations suggest that the first line of contact between Env and an antibody is through a loop region(Schmidt et al., 2013). Accordingly, for our analysis, we have used a probe size of 6Å, which is the typical diameter of a hairpin loop. At each residue present on the surface of the protein, the approaching probe was considered in three directions (**Figure 6A**): perpendicular to the surface **z**, and then the **x** and **y** directions spanning the plane parallel to the surface. The geometric mean of the **x**, **y** and **z** values provide the GEF per residue on the Env surface. This value will go to zero when the glycan encounter factor is zero from any one of the three cardinal directions. Thus, for any point on the surface which has a dense glycan covering, such as D in **Figure 6B**, has a high glycan encounter factor value, versus a point such as R where the glycan covering is sparse, which will have a low GEF.

**Figure 6:**
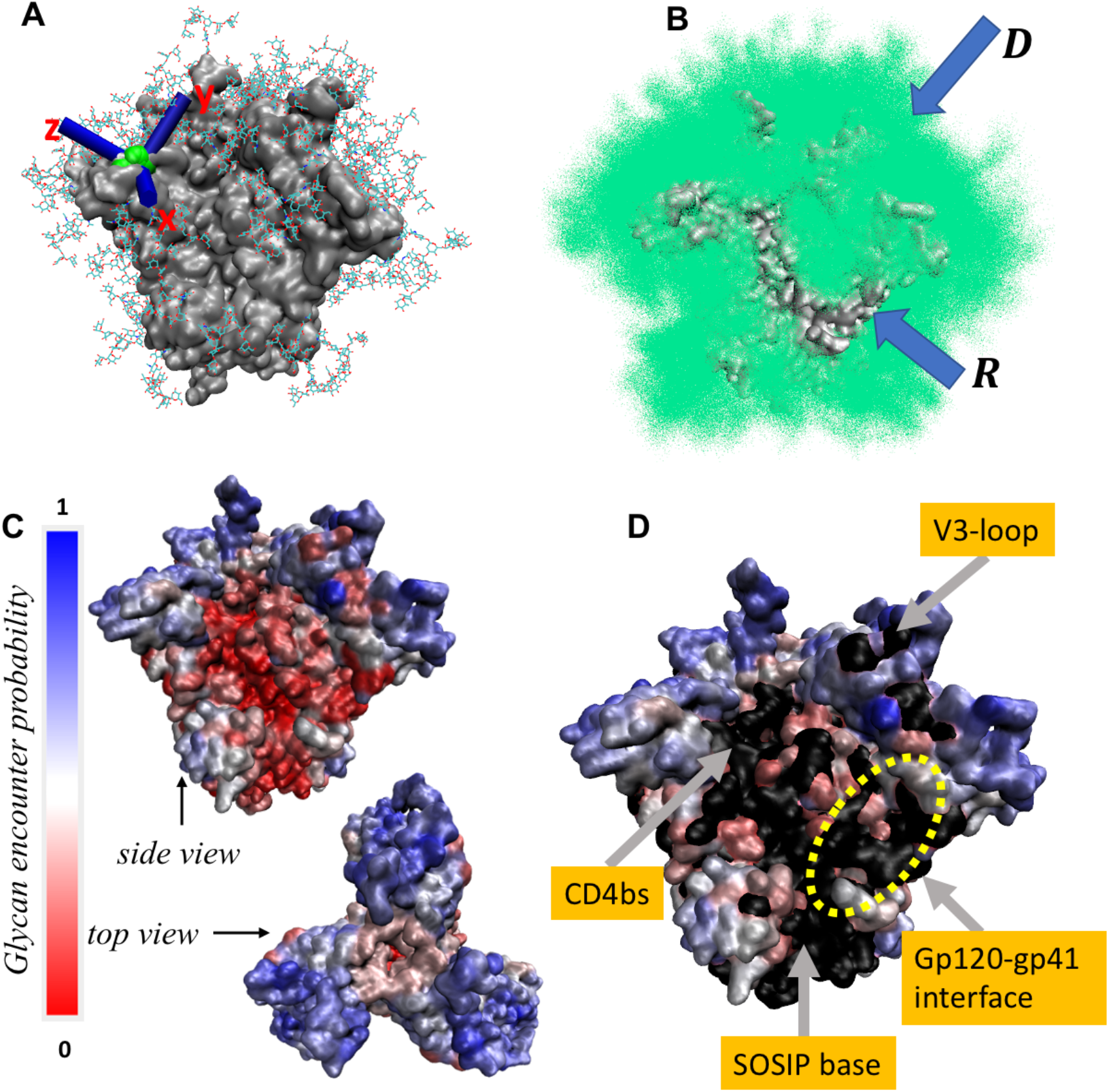
Glycan Encounter Factor (GEF) for quantifying shielding effect. (A) At each residue present on the surface of the protein, the approaching probe is considered in three directions **x, y** and **z**. (B) Any point on the surface which has a dense glycan covering, such as D, has a high glycan encounter factor value, versus a point such as R where the glycan covering is sparse, which will have a low GEF. (C) Representation of GEF on Env surface, given by a colormap. Regions of BG505 surface having a GEF less than 0.14 is colored in black (D). Typical known antibody epitopes are indicated by arrows. BG505-specific GH and COT epitope region demarcated by yellow dashed circle.

Previous evidence suggests that 70% of the Env ECD surface area is covered by glycans(Pancera et al., 2014). Based on this, we determined the lower cut-off of GEF below which we can define glycan holes. **Figure 6C** shows GEFs mapped onto the trimer protein surface. The calculated solvent-exposed surface area of the protein part of our modeled BG505 structure without considering glycans is 86,055 Å^2^. Excluding the exposed region at the base of the soluble SOSIP, this reduces to 79,672 Å^2^. Varying the lower cut-off of GEF, we find that a cut-off of 1.5 GEF leads to 30% of the surface to be exposed. Regions of BG505 surface having a GEF less than 1.5 is colored in black in **Figure 6D**. From the figure, it is clear that the typical glycan holes targeted by bNAbs in BG505, such as the CD4 binding site, the V3-loop epitope and the fusion peptide binding region fall below this GEF cut-off. GEF tracks with epitopes that are relatively generic to a broad range of Env strains(Burton et al., 2012; Xu et al., 2018).

Antibodies elicited by BG505.SOSIP.664 are mainly biased towards the missing 241 and 289 glycan holes (GH) and the cleft-of-trimer (COT) epitope regions as demonstrated by cryo-EM and immunogenicity assays(Bianchi et al., 2018; McCoy et al., 2016) (see Figure 1 in reference (Bianchi et al., 2018)). The GEF parameter identifies these BG505-specific epitopes as breaches in the glycan shield (**Figure 6B**, “R” region; **Figure 6D** yellow dashed circle). On the other hand, the densely glycosylated regions around V2, V4 loop and alpha 2 helix have high values of GEF. At each point, the GEF value is given by a combination of all glycans in the vicinity that can come in the way of the approaching probe. At the same time, we can analyze the extent of influence of each glycan on the protein surface. Thus, we are now equipped with a tool to quantify the barrier effect of the glycans individually or as a group for immunogen design, as further demonstrated(Bianchi et al., 2018; McCoy et al., 2016) in the following section.

### Combining network topology and glycan encounter factor to inform on local and global effects of neutralization

The proposed network is useful in deciphering the impact of addition or deletion of glycan on neutralization. Removal of the high conserved glycan at 197 by mutating the sequon leads to enhanced neutralization sensitivity to a variety of CD4bs and V3-specific antibodies(Huang et al., 2012; Li et al., 2008; Townsley et al., 2016). This glycan is situated proximal to the CD4 binding site, and the tip of V3 loop, directly affecting the binding of these antibodies. However, past experimental evidence also suggests that the deletion of this glycan at N197 increases the binding affinity of two antibodies PG9 and PGT145 (Behrens et al., 2016) which target the trimer apex of Env. The epitope of PG9 as determined from the PDB structure 5VJ6 include residues 160, 161, 167-173, 185, 305 and 307 (**Figure 7A**) and the epitope of PGT145 as determined from PDB structure 5V8L include residues 123, 124, 127, 160-162, and 167-169 (**Figure 7A**). The footprint of glycan 197 as per our GEF model is shown in **Figure 7B**. Residues of the antibody epitopes do not overlap with those regions directly covered by glycan 197. Yet, removal of glycan 197 results in significant reduction in the glycan encounter factor over the V2 antibody epitope as evidenced in **Figure 7C** and **D**.

**Figure 7:**
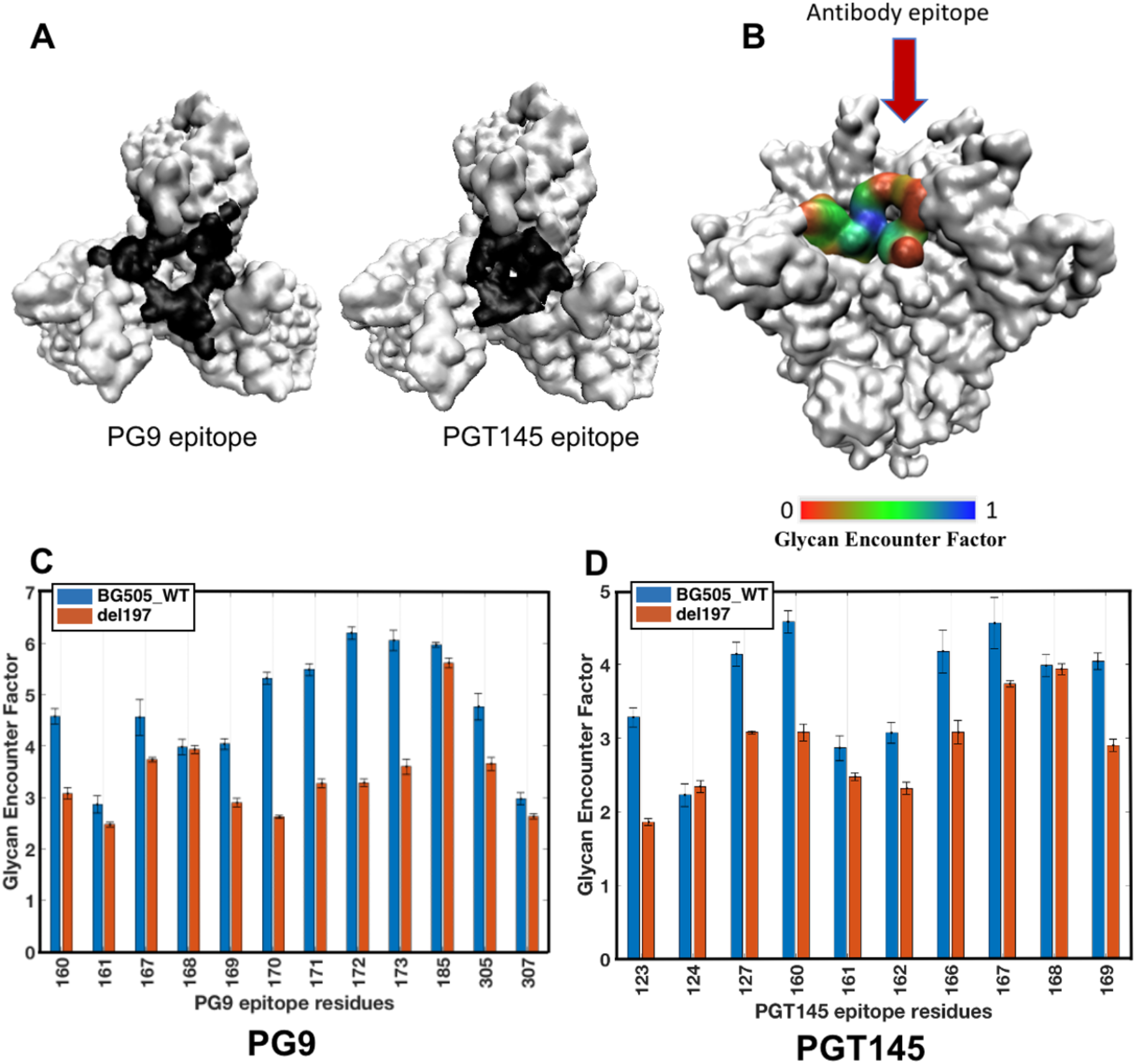
Deletion of glycan N197 decreases glycan shielding at PG9 and PGT145 epitope. (A) Top view of Env showing epitope regions of antibodies PG9 and PGT145 at the apex. (B) Footprint of glycan N197 on Env surface, colored by glycan encounter factor contributed by N197 alone. (C) Glycan encounter factor over the PG9 and (D) PGT145 epitope residues. GEF decreases for both the epitopes due to deletion of N197 glycan.

The removal of glycan 197 affects the glycans that were originally acting as barriers over the epitopes to resample the available space in such a way that the barrier over the epitopes is now reduced. Glycans 156, 160, 185e and 185h from the neighboring protomers directly shield the PG9 and PGT145 epitopes. Looking at the shortest paths of communication between residue 197 from either of the protomers to all other glycans (**SI Figure S6A, B**) demonstrate that while the shortest path between glycans 160 and 197 is direct, other glycans covering the epitope regions communicate with 197 across neighboring protomers or via a series of inter-glycan interactions. Deletion of 197 does not only affect the V1/V2 loop region glycans. The difference in adjacency matrix due to this perturbation is illustrated in **SI Figure S7A**. There is a slight modification of the community structure within the network as well, with glycan 392 leaving module 4 (red) and entering module 3 (yellow) instead (**SI Figure S7B**). Interactions such as those between glycans N88 – N625, N133 – N363, and interprotomer N156 - N185e are reduced more than 10%. On the other hand, glycan 276, which was originally interacting strongly with 197 now forms new interactions with N295, N332 and N411 glycans (**SI Figure S7C, D**).

## Discussion

Dense arrangement of N-glycans masking antigenic surfaces on Env acts as a dynamic shield from the adaptive immune system. Understanding the structure and dynamics of the glycan shield as a whole is therefore important for characterizing the immunologically inert regions of Env. While high-resolution X-ray, NMR and cryo-EM structures have supplied a number of important molecular details about the Env glycoprotein, they do not account for the high flexibility and dynamics of these glycans that leads to the glycan shield. Our integrated method for capturing dynamics of the glycans beyond the core β-mannose, and quantification of the barrier resilience and vulnerability as a function of glycosylation pattern can add new perspective to current HIV vaccine design efforts.

Previous computational studies have generalized glycan moieties as simple oligomannoses such as mannose-5 and mannose-9 (Ferreira et al., 2018; Lemmin et al., 2017; Liang et al., 2016; Stewart-Jones et al., 2016; Yang et al., 2017) for the purpose of modeling. For the first time, we have incorporated native glycosylation, and included complex sugars based on site-specific mass spectrometry results. While the overall aspects of the network are consistent between mannose-9 and native-like glycosylation, there are critical differences. The most noteworthy are a consequence of the presence of the bulky fucose ring at the base or the negatively charged sialic acids at the antennae tips. These lead to overall rearrangement of glycan orientations affecting its microenvironment, ultimately influencing the shield topology. Thus, our comprehensive characterization of the shield is capable of capturing individual glycan effects which are physiologically and immunologically more relevant.

Our network-based approach illuminates the collective behavior of glycans. We compare the relative centrality of glycans, identify potential interaction pathways and detect stable communities. The centrality or importance of glycans correlates well with experimental cryo-EM data(Amanat et al., 2018; Berndsen et al., 2019). Glycans with lower centrality have lesser influence on the graph, and are the first to be eliminated from the network, if the adjacency threshold is increased. Similarly, the most central region of the network is the most resilient to enzymatic action. Such centrality measures can help determine the ease of targeting glycans and their modifications, guiding the process of immunogen development in the context of distinct Envs. While the shield resilience is the main consideration for epitope exposure, some antibodies have also evolved to capitalize on specific interactions with a number of the conserved glycans during their molecular engagement with the epitope. Due to the stability of glycan interactions within the detected communities, this help determine the antibody angle of approach that has been shown to influence the breadth of bNAbs (Moore and Williamson, 2016).

There are known common and uncommon ‘glycan holes’ that can open up in the shield to make the virus more vulnerable, or conversely get covered resulting in immune escape, as a result of evolutionary addition/deletion or shift of glycans(Wagh et al., 2018). While some advances have been made previously in identifying these breaches based on the area of influence of each glycan(Wagh et al., 2018), here we have found that this area can vary, depending on the glycan type, charge, neighbors, etc. Amino acid signature analyses suggest that even minor perturbations such as single site mutations could potentially change the shielding effect over certain epitopes(Bricault et al., 2019). Therefore, we have derived a measure to quantify the shielding effect based on the encounter of glycans over the antigenic protein surface. This tool allows us to define regions of relative vulnerability and resilience in the glycan shield.

The method we developed here for the structural modeling of the glycoprotein atomistic ensemble and the subsequent development of the network is high-throughput compared to traditional sampling by MD simulations. Due to the ease of this fast and efficient pipeline we are now equipped with a tool to perform comparative structural studies due to glycan additions, deletions or modifications, as well as other variations in the Env sequence. This is important to understand the evolution of the glycan shield over longitudinal sampling of lineages. The structural basis of addition or removal of glycans that are known to drive antibody maturation and neutralization activities(LaBranche et al., 2018; Umotoy et al., 2019) can also be easily investigated. Capitalizing on the relatively low computational overhead, our pipeline can be integrated into a polyclonal epitope mapping assay to track the glycan shield as a function of hierarchical antibody response, by modeling the shield network with the presence of different antibodies and observing how the topology is altered.

Beyond HIV-1 Env and other viral envelope proteins, the significance of glycoproteins in a vast array of biological processes from protein folding, cellular communication to immune-regulation, make them a fast-emerging field of interest in biomedical research. Changes in these glycosylation patterns have been associated with various diseases, including rheumatoid arthritis(Nakagawa et al., 2007), and cancers(Taniguchi and Kizuka, 2015). Additionally, many of the current therapeutic antibodies in the market are N-linked glycoproteins, and the significance of N-glycans is becoming increasingly evident (Dalziel et al., 2014). The ease of modeling of glycan network utilizing our approach makes it translatable to other systems, and can assist in determining the role of these glycans in conjugation with the underlying proteins at a molecular level. In comparison to N-glycans, O-linked glycosylations are relatively more complex in terms of structural variations and our understanding is limited. Generalization our methodology to encompass O-glycosylated systems can provide molecular insights on such refractory systems, like Ebola that have both N- and O-linked glycosylations, and explore the pathological implications of dense O-glycans in mucin associated cancers(Bhatia et al., 2019).

In summary, HIV and the human immune system are at war constantly, and the virus uses the Env glycan shield to mask the human immune surveillance. One battlefront in this war is the glycan shield: while the virus evolves to develop resilience, the immune response counteracts by looking for vulnerabilities. As efforts are underway to aid the immune system overcome this race by conditioning it with engineered immunogens, there is a need to quantify the resistance and vulnerability of glycan shield in a more quantitative manner. This is that first time that the native glycan network and shielding has been spatially quantified. Our derivation of the Glycan

Encounter Factor measures the relative barrier over the Env protein surface, and can potentially aid to distinguish subtle differences on the shield due to variations in the glycosylation or even protein sequences. We are therefore armed with a set of validated *in silico* tools with which to help the anti-HIV war efforts and guide immunogen design.

## STAR Methods

### Contact for Reagent and Resource Sharing

Further information and requests for resources and reagents should be directed to and will be fulfilled by the Lead Contacts S. Gnanakaran and Andrew B. Ward.

### Method Details

#### High-throughput conformational modeling

The ensemble of BG505 glycoprotein 3D conformations were built in atomistic detail by implementing the ALLOSMOD(Guttman et al., 2013; Weinkam et al., 2012) package of MODELLER(Eswar et al., 2006; Sali, 1995) [RRID: SCR_008395]. For ease of understanding, the modeling protocol has been described as a flow schematic in **SI figure S8**. The protocol consists of the following steps:

a. ***Base template selection for protein scaffold:*** The BG505 protein scaffold was homology modeled, by threading the protein sequence into available crystal structure templates. Ligand-free SOSIP structure with PDB accession ID 4ZMJ(Kwon et al., 2015) was used as the base template for gp120, that also guided the three-fold symmetry of the trimer. 5CEZ(Garces et al., 2015) was used as the base template for gp41 and since it has the least number of missing residues among the available structures.
b. ***Identifying flexible regions from available experimental PDB structures to guide conformational sampling:*** There are a large number of unstructured flexible regions in the protein. In order to incorporate the fluctuations as dictated by the available structural data we devised the following protocol. We aimed to construct 10 distinct protein models for these flexible regions as starting scaffolds for glycan building. In order to model these, 34 different BG505 SOSIP experimental structures were selected below a resolution cut-off of 4Å (see **SI Table S2**). Open-like, chimeric and synthetic models were not considered. The set of 34 were structurally aligned using PyMOL(Schrödinger, 2015) and their residue-wise backbone RMSF was calculated (**SI Figure S9A**). The base-level of 0.1 was used as a threshold in the normalized RMSF measure. The unstructured residues (coils and turns) were identified on 4ZMJ and 5CEZ using the STRIDE program(Frishman and Argos, 1995). All the identified regions with RMSF higher than the threshold (**SI Figure S9A**) were considered for modeling based on multiple templates. These include residue ranges 64 – 82, 133 – 153, 184 – 189, 298 – 303, 312 – 314, 324 – 329, 396 – 413, 425 – 431, 457 – 464, and 609 – 618 (in HXB2 numbering).
c. ***Template selection for identified flexible regions:*** Some of these identified regions have missing residues in many of the PDBs (**SI Figure S9B**). Wherever enough models were available, 10 distinct conformations were randomly selected as templates. If this number 10 could not be reached for some regions (for example, in loops V2 and V4) due to missing residues (**SI Figure S9B**), those residues were modeled *ab initio.* All known disulphide bonds were kept as additional restraints. We made sure these short-listed regions for *ab initio* modeling were less than MODELLER-prescribed 14 residues in length. This is because, modeling of such long loops are usually considered to be error-prone(Fiser et al., 2000). The loop region 546-569 in gp41 is present only in the structure 5CEZ. We did not model this region *ab initio*, and used 5CEZ as the only template for all models. The 10 models thus built were further tested for proper stereochemistry, based on MODELLER optimization scores and PROCHECK scores(Laskowski et al., 1993). This results in different starting orientations of the hypervariable loops, within the experimentally determined limits of fluctuations.
d. ***Modeling of glycans and their ensemble sampling:*** For each of the 10 selected protein structures, glycans were initially added with random orientation, at the known glycosylation sites, based on ideal geometries as dictated by CHARMM36(Best et al., 2012; Huang and MacKerell Jr, 2013) force field internal coordinates, followed by a 1Å randomization added to the overall atomic coordinates as described by Guttman et. al.(Guttman et al., 2013). Once all the glycans were added, ensuing refinement steps of the glycoprotein system optimized an energy function given by a combination of template-based spatial restraints, CHARMM36 forcefield terms, and a soft sphere-like truncated Gaussian term to prevent collisions. The structures were relaxed with 1000 steps of conjugate gradient minimization followed by a short molecular dynamics equilibration of 500ps. Further refining with five rounds of simulated annealing was performed between 1,300K to 300K in 8 steps. The glycans and the loop regions were kept flexible during the refinement steps. 100 fully glycosylated structures were modeled from each of the 10 selected protein models, resulting in the final ensemble containing 1000 different poses. Template restraints were removed from the hypervariable regions with glycans in order for them to sample a wider range of conformations during the glycoprotein relaxation phase. These residues by HXB2 numbering are as follows: 143 to 152 (V1 loop), BG505-specific insert residues 185A to 185I and 186 to 189 (V2 loop), 309 to 315 and 325 to 329 (V3 loop), 400 to 410 (V4 loop) and 458 to 464 (V5 loop) as determined from the LANL HIV database (https://www.hiv.lanl.gov/).

#### Glycan root mean square fluctuations and volume calculation

For both the native and all-man9 glycosylated models, the root mean square fluctuations (RMSF) of each glycan (with index *n*) was calculated as an average over all its heavy atoms, by the following equation:

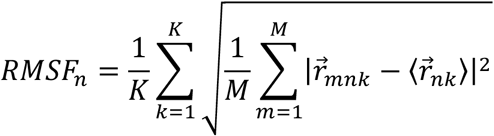

where 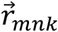 is the atomic position of heavy atom *k* of glycan *n* in snapshot *m*, 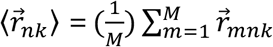 is the average atomic position of heavy atom *k* in glycan *n*. K is the total number of heavy atoms in the glycan. It is 127 for mannose-9, and varies depending on the type of glycan. The ensemble for each model contains 1000 snapshots, making *M* = 1000 snapshots for each of the two models. The standard deviations were obtained by dividing the 1000 snapshots into 4 sets of M=250, and calculating the four sets of RMSF values (**SI Figure S5**).

The 1000 structures of the total ensemble are built from 10 initial starting protein conformations, as described above. The main difference between these 10 conformations are the variations in the loop regions, due to the missing residues in the templates. In order to remove the effects of the loop fluctuations and consider the RMSF coming from the glycans alone, the reduced RMSF values were also calculated in each of these 10 sub-models, and their average and s.d. calculated (**SI Figure S5B**). The RMSF difference between the models (**SI Figure S5E**) were obtained by subtracting the reduced RMSF values of all-man9 from native model (native minus all-man9). Since, the average reduced RMSF value is ~4Å, only those RMSF difference values are statistically significant which are above 0.2 Å, which corresponds to a p-value of 0.05, rejecting those values within the null hypothesis. Ensemble of individual glycans were converted to atomic densities at 1 Å gridsize in order to calculate the volume. Volume calculation was done using CHIMERA package(Pettersen et al., 2004).

#### Glycan overlap and network analysis

The inter-glycan overlap is calculated as the total fraction of heavy atoms from the two glycans that come within 5Å of each other. Let us consider the example of mannose-9 to illustrate the parameter of overlap. A single mannose-9 glycan has 127 heavy atoms. Since our ensemble is composed of 1000 possible structures, there are effectively 127,000 heavy atoms per ensemble of mannose-9 at one position. The fraction of the total number of heavy atoms from two neighboring ensembles that come within contact distance defines the overlap fraction. Since mannose-9 is the most commonly occurring glycoform in our system, we have used it as our reference for normalization of the overlap probability. An overlap greater than or equal to 50% of heavy atoms from two neighboring mannose-9 glycans is assigned as 1. This overlap matrix is used to define the adjacency matrix for our network analysis. Each glycan functions as a node of the graph (**Figure 4A inset**), and two nodes are connected by an edge if there is at least 5% overlap as per our overlap definition given above. The edge length is inversely proportional to the overlap value, i.e., the larger the overlap, the closer two nodes (glycans) are in the graph. Only those glycans from the neighboring protomers are considered, that have an inter-protomer edge. All graph theory and network analyses were performed using Python(Rossum, 1995) [RRID: SCR_008394] and Matlab_R2018a packages(MATLAB, 2018) [RRID: SCR_001622].

#### Eigenvector centrality calculation

For a given graph, *G*, with adjacency matrix **A**=(*a_v,t_*) where *a_v,t_* is the edge weight connecting nodes *v* and *t* (*a_v,t_* = 0 when there is no connection), the relative centrality score *x*, of node *v* can be defined as:

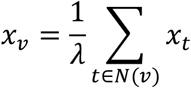

where *N*(*v*) is the set of neighbors directly connected to *v*, and λ is a constant. From the definition of the adjacency matrix where the elements go to zero if two nodes are not connected, the above equation can be expressed as:

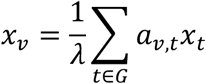

This has the form of the eigenvector equation **Ax**= λ**x**. With the added constraint of the eigencentrality values needing to be non-negative, by the Perron-Frobenius theorem(Pillai et al., 2005), the eigenvector corresponding to the largest eigenvalue gives the desired measure of centrality. The eigenvector is a unit vector and therefore the centrality values add up to one. For the purpose of this work, we have normalized the centrality values with respect to the node with the highest centrality assigned at 1, to obtain the relative centrality values.

#### Modularity maximization for community detection

Community detection within the glycan network was performed using the modularity maximization approach given by Newmann and Girvan(Newman, 2006; Newman and Girvan, 2004). Modularity Q is calculated as the difference between the fraction of edges that fall within a module and the expected fraction if the edges were distributed in random.

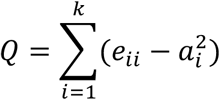

Where *e_ii_* is the fraction of edges in module *i*; *a_i_*, is the fraction of edges with at least one end in module *i*. It is calculated by a greedy heuristic, beginning with the trivial system of each node being a cluster, and merging two clusters that will increase modularity by the largest value, stopping when any further merge would decrease the modularity. This approach is known to work well for small networks similar to our system. The calculations were implemented through a standard algorithm(Blondel et al., 2008) in Matlab R2018a.

##### Visualization of Structures

All visualizations of structures for figures were generated using the Visual Molecular Dynamics package(Humphrey et al., 1996) [RRID: SCR_001820].

## Data Availability

All data needed to evaluate the conclusions in the paper are present in the paper and/or the Supplementary Materials, and further deposited in the library: http://dx.doi.org/10.17632/yds8rtfjjy.1

Additional data related to this paper may be requested from the authors.

## Supplemental Information

Supplemental Information includes details of torsion angle distributions, flexibility and volume sampling analysis of modeled glycan structures, along with 2 Tables and 9 Figures.

## Acknowledgements

S.C., B.T.K and S.G were funded by grants from the NIH (Center for HIV/AIDS Vaccine Immunology and immunogen Discovery, UM1 AI100645; Consortia for HIV/AIDS Vaccine Development, UM1 AI144371). Z.T.W and A.B.W were supported by the National Institute of Allergy and Infectious Diseases grants: UM1 AI100663 and UM1 AI144462, and Collaboration for AIDS Vaccine Discovery (CAVD) grant OPP1115782. S.C was also partially supported by the Center for Nonlinear Studies (CNLS) at Los Alamos National Laboratory (LANL). We acknowledge Timothy Travers and Cesar Lopez for suggestions on glycan modeling, Kshitij Wagh for insights on glycosylation holes in HIV, Kevin Weihe and Rory Henderson from Duke University for helpful discussions on Env immune response. S.C and S.G acknowledge the LANL High Performance Computing Division for providing computational facilities.

## Author Contributions

Conceptualization, S.C., Z.T.B, N.W.H., B.T.K., A.B.W. and S.G; Methodology, S.C., Z.T.B, N.W.H., A.B.W. and S.G.; Modeling and Simulations, S.C.; Cryo-EM experiments, Z.T.B.; Formal Analysis, S.C. and Z.T.B.; Visualization, S.C. and Z.T.B.; Manuscript Preparation, S.C., Z.T.B, N.W.H., B.T.K., A.B.W. and S.G.; Supervision, A.B.W. and S.G.

