## Supplemental Information for "Quantification of the Resilience and Vulnerability of HIV-1 Native Glycan Shield at Atomistic Detail"

**Modeled glycan ensembles sample a biologically relevant landscape:**

The PNGS asparagine chi1 dihedral, and the phi and psi dihedral distributions of 9 different inter-glycan linkages within the ensemble was compared with those obtained from different glycan structures available in the PDB database, as described in **Figure S1**. Torsion angle distributions from the PDB were obtained using GlyTorsion (Lütteke et al., 2005). These distributions match well between our generated ensemble and the PDB structures, Four of the psi angles, namely, the (i)1-3 linkage between two α-mannose sugars; (ii) 1-3 linkage between α-mannose and β-mannose; (iii) 1-2 linkage between two α-mannoses; and (iv) N-acetylglucosamine and α-mannose sample a wider distribution of angles in our models. It must be kept in mind that while the PDB distributions included ~17,000 structures for the ASN and ~7,000 structures for 1-4 linkage between two N-acetylglucosamines at the glycan core, there were less than 2000 structures for all other distributions. Also, these sugar linkages in the PDB come from both N-glycan and O-glycan types, have conformational restraints stemming from experimental structure determination, and are not nearly as densely packed as in our HIV model. Our ensemble, on the other hand, has 84 glycans, with 1000 conformations each. The larger number of structures per linkage, as well as the dense interactions between neighboring glycans can result in some of the distributions sampling wider range of torsion angles.

**Glycan fluctuations and their dependence on complex sugars**

Glycan dynamics, type, and inter-glycan interactions determine the local shielding effect over the Env protein. We have looked at the dynamics of individual glycans and how they change due to native-like glycosylation, ultimately governing the glycan shield properties. Each glycan samples a large region in space as shown in **Figure S3A and S3B**. These fluctuations of course become much more extensive, when the glycans are present on the variable loops, due to the dynamic movement of the protein backbone of the loops at these regions. The root mean squared fluctuations (RMSF) measure and the sampled volume for those glycans located on the loops are generally much higher, as seen in **Figure S5A** and **Figure S5C**. However, aligning the protein backbone and considering the reduced RMSF contribution coming only from the glycans (see Methods), we see that the fluctuations between different glycans are comparable, with a maximum difference of ~1.5Å (**Figure S5B**). As for the glycan specific sampled volume, after removing the flexibilities brought on by protein loop motions, the glycans on loop V2 and V4 and those in gp41 cover the largest regions in space (**Figure S5D**).

Considering the differences between native-like glycosylation and the all-man9 model, some interesting consistent patterns emerge. Comparing the reduced RMSF between native and all-man9 glycosylation in **Figure S5E,** we see that while glycans 185h, 197, 355, 398 and 406 have decreased fluctuations in the native model, those of glycans 88, 462, 611, 618, 625 and 637 have increased significantly. With the exception of 355, which is a fucosylated hybrid (FH), all of these glycans are complex sugars. The presence of the charged sialic acid at the tips of these complex glycans can dictate the interactions between neighbors, increasing their structural variations if surrounded by other charged glycans, or reducing them if a stable conformation buried between uncharged high-mannose patches can be found. The FH glycan at N355 itself is centrally located between a number charged glycans, those at 88, 398, 462, and some of the gp41 sugars. The reduced fluctuations of this glycan can stem from its attempt to screen these neighboring negatively charged sugars, thus forming stabilizing interactions. The complex sugars in gp41 (611, 618, 625 and 637) have also been shown to have high variations in cryo-EM maps(Ward and Wilson, 2017). The total volume sampled is generally larger for native glycosylation, compared to the all-man9 model (**Figure S5F)**. This is not unexpected, since complex glycans generally have larger number of sugars, including the bulky fucose ring at the base. The centrally located high-mannose patch has almost similar volumes in both the models.

**Table S1:** Different glycan species at BG505 SOSIP PNGS selected based on past site-specific mass spectroscopy data. Structures corresponding to each species is given in Figure 1A (main text).

| **Glycan ID** | 88 | 133 | 137 | 156 | 160 | 185e | 185h | 197 | 234 | 262 | 276 | 295 | 301 | 332 |
|---|---|---|---|---|---|---|---|---|---|---|---|---|---|---|
| **Glycan type** | FA2 | M9 | FA2 | M9 | M8 | FA2 | FA2 | FA2 | M9 | M9 | M7 | M9 | M9 | M9 |

| **Glycan ID** | 339 | 355 | 363 | 386 | 392 | 398 | 406 | 411 | 448 | 462 | 611 | 618 | 625 | 637 |
|---|---|---|---|---|---|---|---|---|---|---|---|---|---|---|
| **Glycan type** | M9 | FH | M9 | M9 | M9 | FA2 | FA2 | M9 | M9 | FA3 | FA3 | FA3 | FA2 | FA2 |

**Table S2:** List of experimental BG505 SOSIP structures used for flexibility analysis. PDB accession IDs are given. BG505_293S and 293F are cryo-EM structures with high oligomannose and native-like glycosylated structures obtained in our previous work(Berndsen et al., 2019). 5CJX has three protomers having different loop orientations. Therefore, they were considered separately.

| Structure | Resolution (Å) | Method |
|---|---|---|
| 6MTJ | 2.34 | X-Ray |
| 6MTN | 2.5 | X-Ray |
| 6MU7 | 2.5 | X-Ray |
| 6MU6 | 2.55 | X-Ray |
| 6NNJ | 2.6 | X-Ray |
| 6NM6 | 2.74 | X-Ray |
| 6NNF | 2.76 | X-Ray |
| 5V7J | 2.91 | X-Ray |
| 6MU8 | 2.99 | X-Ray |
| 5CEZ | 3.03 | X-Ray |
| 5U7M | 3.03 | X-Ray |
| 5U7O | 3.03 | X-Ray |
| 4TVP | 3.1 | X-Ray |
| 5FYL | 3.1 | X-Ray |
| BG505_293S | 3.1 | Cryo-EM |
| BG505_293F | 3.1 | Cryo-EM |
| 6MPG | 3.2 | Cryo-EM |
| 4ZMJ | 3.31 | X-Ray |
| 6NFC | 3.43 | Cryo-EM |
| 5T3Z | 3.5 | X-Ray |
| 5UTF | 3.5 | X-Ray |
| 6OT1 | 3.5 | Cryo-EM |
| 5CJX_1 | 3.58 | X-Ray |
| 5CJX_2 | 3.58 | X-Ray |
| 5CJX_3 | 3.58 | X-Ray |
| 6CDI | 3.6 | Cryo-EM |
| 6NF2 | 3.7 | Cryo-EM |
| 6NF5 | 3.71 | Cryo-EM |
| 6CDE | 3.8 | Cryo-EM |
| 6CH7 | 3.8 | X-Ray |
| 6MPH | 3.8 | Cryo-EM |
| 6DFH | 3.85 | Cryo-EM |
| 5T3X | 3.9 | X-Ray |
| 6ORO | 3.9 | Cryo-EM |

**Figure S1:** Comparison of PNGS asparagine chi1, as well as phi and psi dihedral distributions of 9 different inter-glycan linkages in the modeled ensemble with those obtained from different glycan structures available in the PDB. Distributions on the left are from the PDB, as given by GlyTorsion. Distributions on the right are from our models. Number of torsions analyzed for each dihedral type from PDB is given.

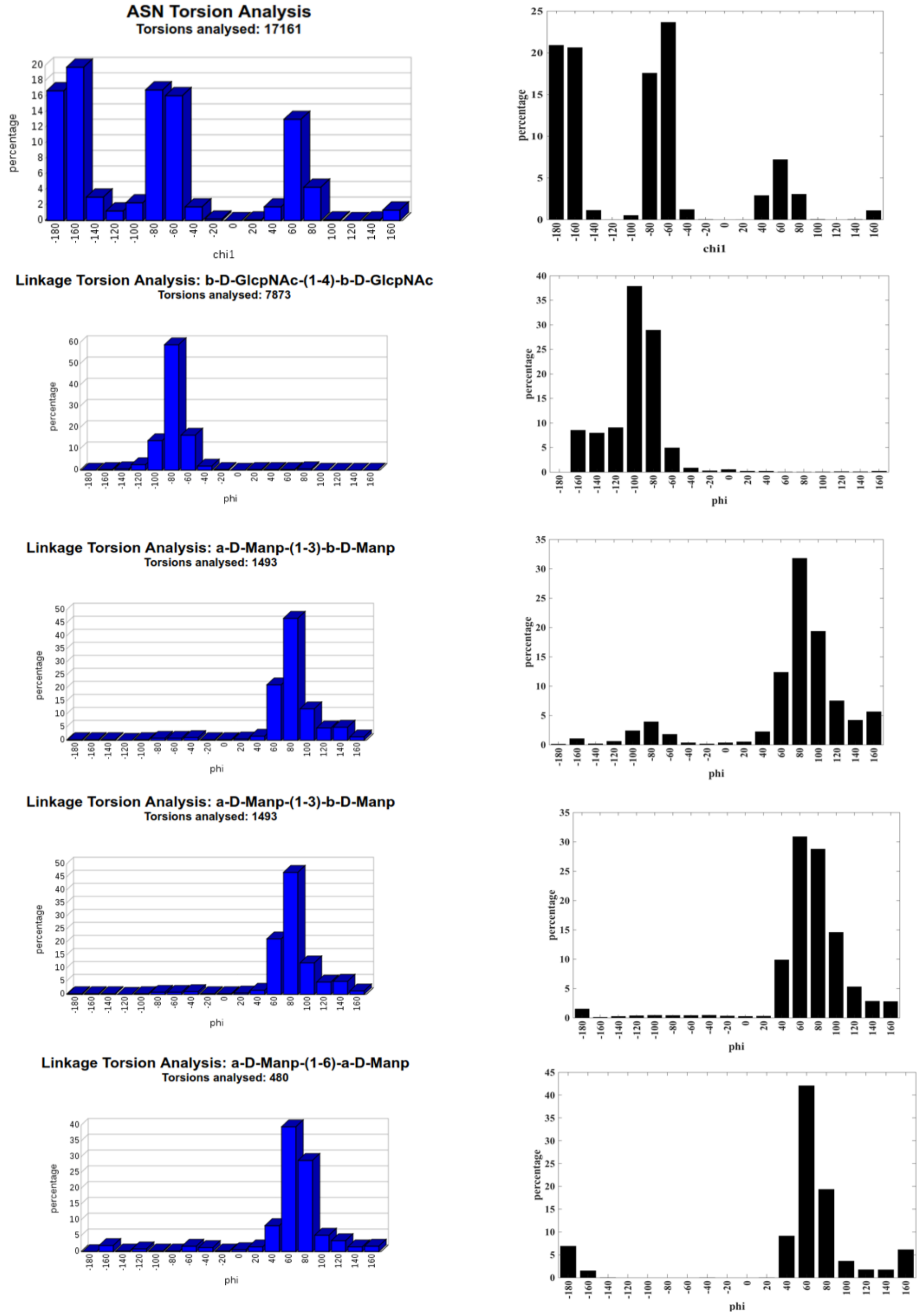

**Linkage Torsion Analysis: a-D-Manp-(1-6)-b-D-Manp**
**Torsions analysed: 1324**

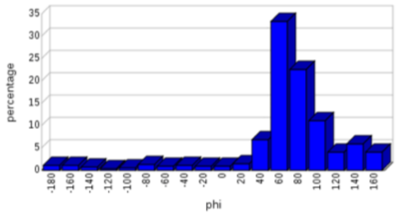

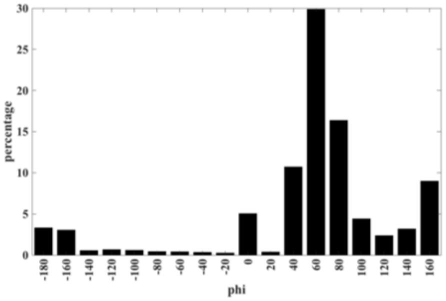

**Linkage Torsion Analysis: a-D-Manp-(1-2)-a-D-Manp**
**Torsions analysed: 819**

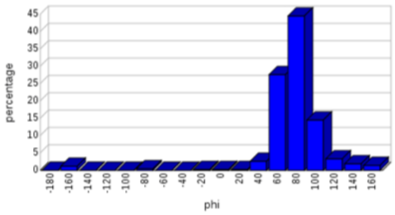

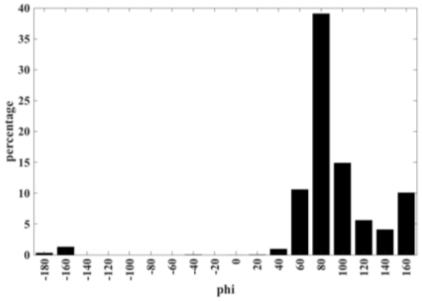

**Linkage Torsion Analysis: b-D-GlcpNAc-(1-2)-a-D-Manp**
**Torsions analysed: 460**

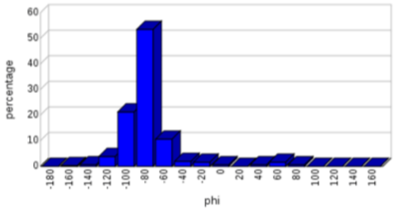

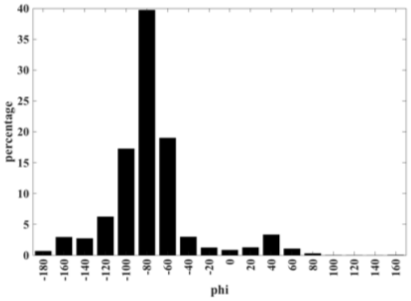

**Linkage Torsion Analysis: b-D-Galp-(1-4)-b-D-GlcpNAc**
**Torsions analysed: 468**

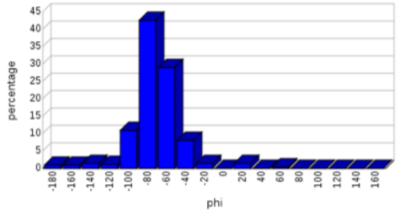

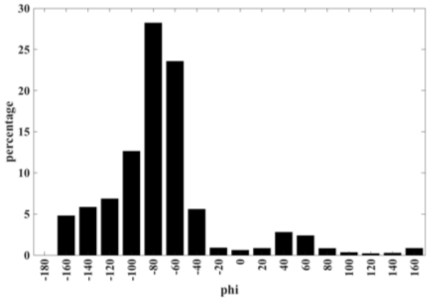

**Linkage Torsion Analysis: a-L-Fucp-(1-6)-b-D-GlcpNAc**
**Torsions analysed: 673**

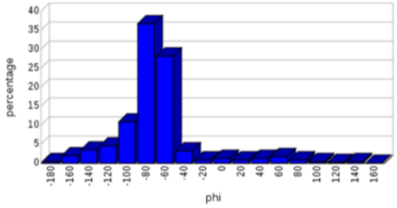

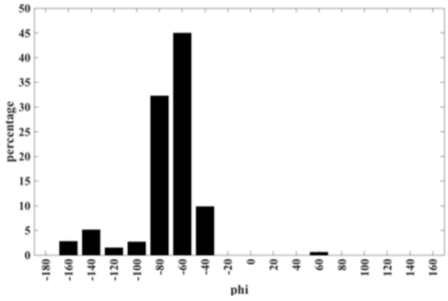

**Linkage Torsion Analysis: b-D-GlcpNAc-(1-4)-b-D-GlcpNAc**
**Torsions analysed: 7873**

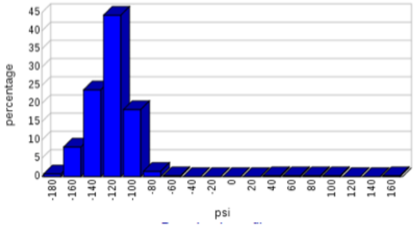

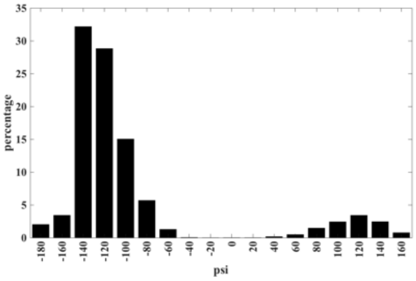

**Linkage Torsion Analysis: b-D-Manp-(1-4)-b-D-GlcpNAc**
**Torsions analysed: 2868**

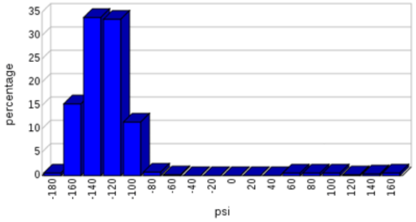

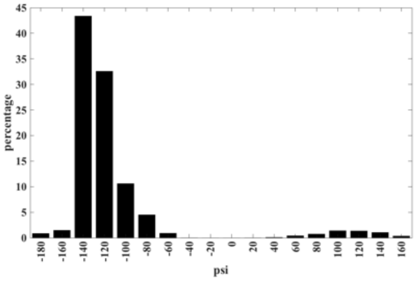

**Linkage Torsion Analysis: a-D-Manp-(1-3)-a-D-Manp**
**Torsions analysed: 536**

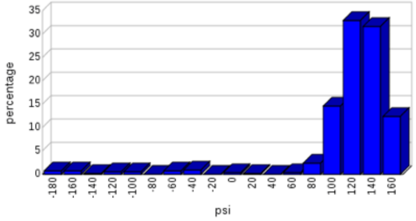

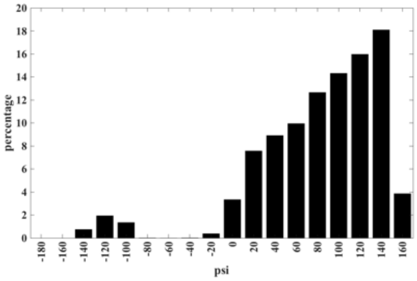

**Linkage Torsion Analysis: a-D-Manp-(1-3)-b-D-Manp**
**Torsions analysed: 1493**

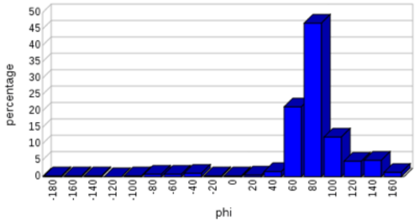

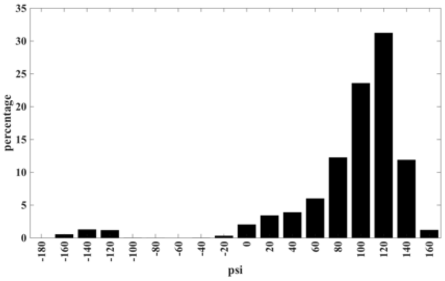

**Linkage Torsion Analysis: a-D-Manp-(1-6)-a-D-Manp**
**Torsions analysed: 480**

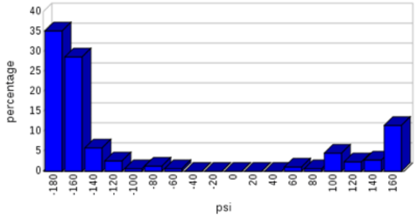

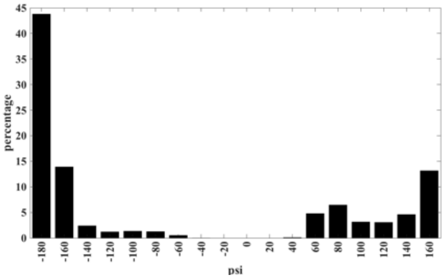

**Linkage Torsion Analysis: a-D-Manp-(1-6)-b-D-Manp**
**Torsions analysed: 1324**

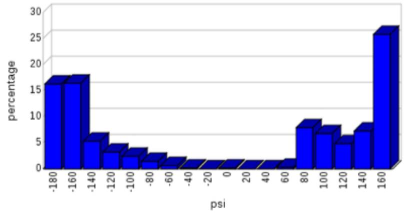

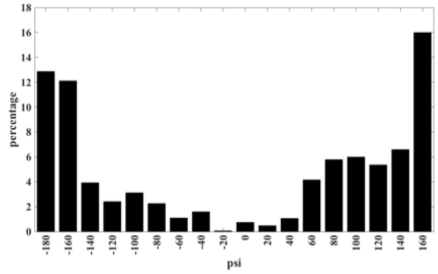

**Linkage Torsion Analysis: a-D-Manp-(1-2)-a-D-Manp**
**Torsions analysed: 819**

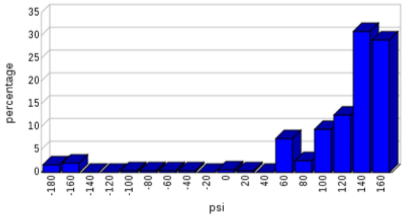

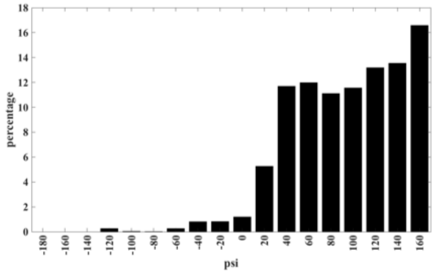

**Linkage Torsion Analysis: b-D-GlcpNAc-(1-2)-a-D-Manp**
**Torsions analysed: 460**

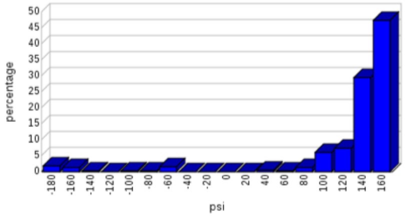

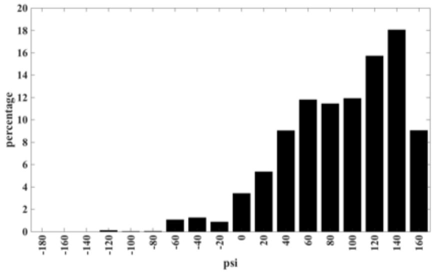

**Linkage Torsion Analysis: b-D-Galp-(1-4)-b-D-GlcpNAc**
**Torsions analysed: 468**

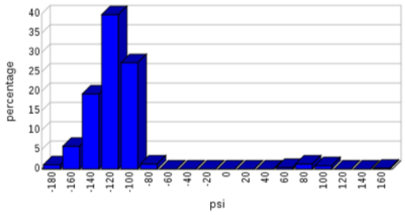

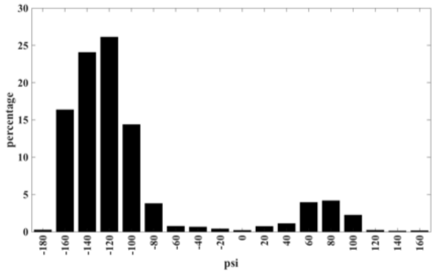

**Linkage Torsion Analysis: a-L-Fucp-(1-6)-b-D-GlcpNAc**
**Torsions analysed: 673**

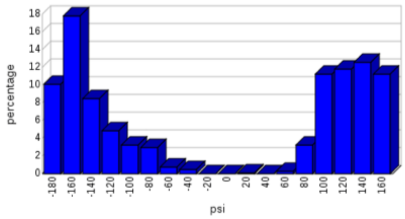

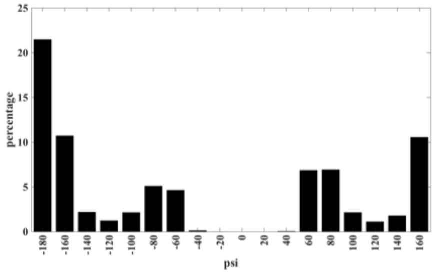

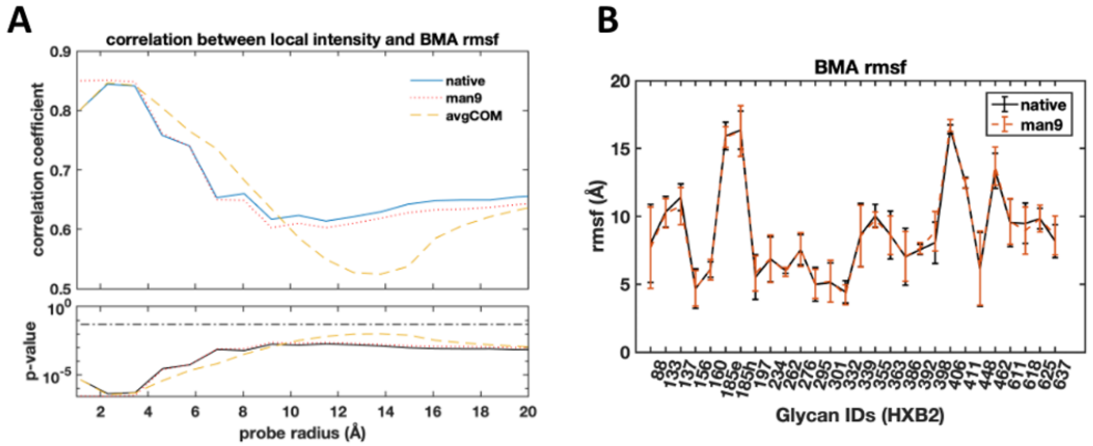

C

| | BG505_293F | BG505_293S | Sim_native | Sim_man9 |
|---|---|---|---|---|
| **BG505_293F** | 1 (0) | 0.90 (1.60E-08) | 0.82 (6.80E-06) | 0.83 (2.30E-06) |
| **BG505_293S** | 0.90 (1.60E-08) | 1 (0) | 0.70 (1.80E-04) | 0.73 (2.30E-04) |
| **Sim_native** | 0.82 (6.80E-06) | 0.70 (1.80E-04) | 1 (0) | 0.95 (1.40E-10) |
| **Sim_man9** | 0.83 (2.30E-06) | 0.73 (2.30E-04) | 0.95 (1.40E-10) | 1 (0) |

**Figure S2: Glycan dynamics are effectively indistinguishable at the level of core BMA between native and all-man9 model** (A) Pearson correlation coefficient and p-value between mean local map intensity around BMA residues and inverse average rmsf of each BMA residue as a function of probe radius. Also plotted is the correlation when using the true C3 average BMA centers of mass (avgCOM). (B) Comparison of BMA root mean squared fluctuations between native and all-man9 models. (C) Table highlighting the correlation coefficients of local BMA intensities between experimental and simulated native and all-man9 maps, p-values given in parentheses. The correlation remains similarly high between all maps at the level of the glycan stem.

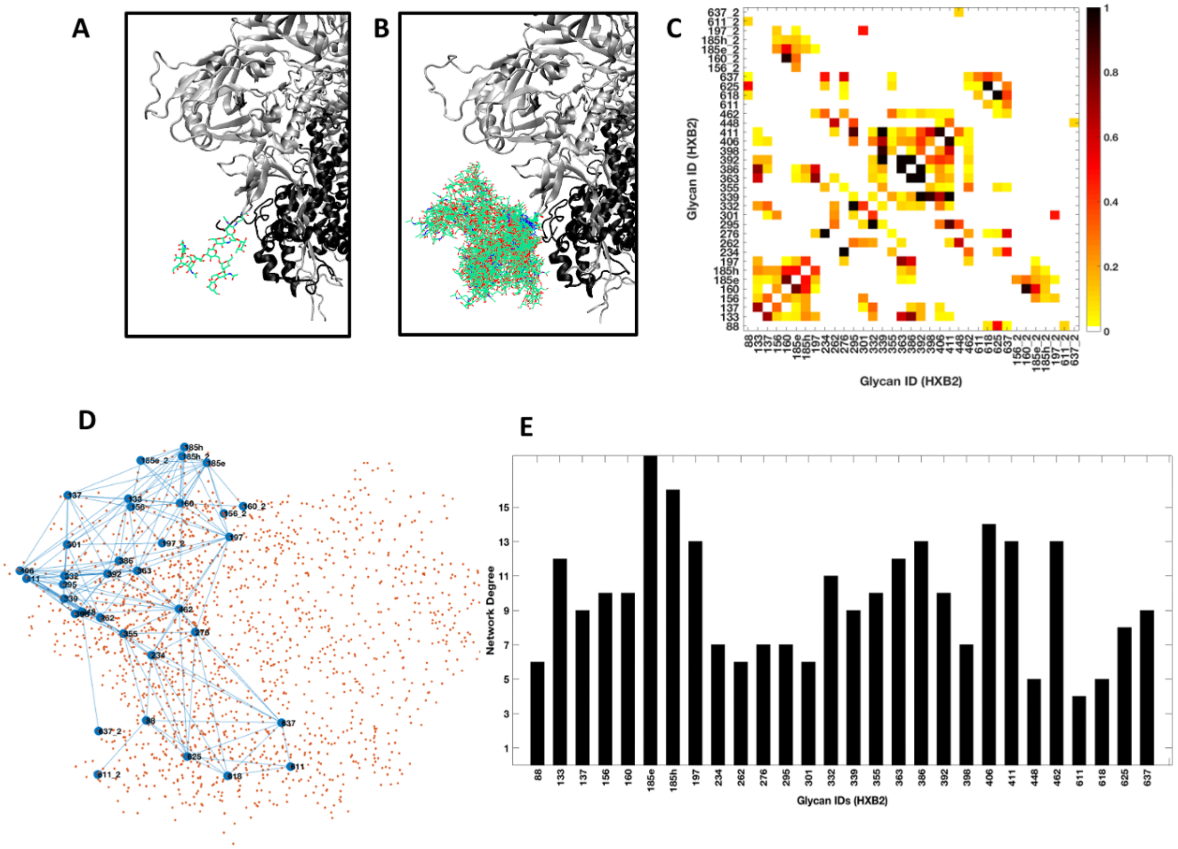

**Figure S3: Topological native glycosylation shield network from glycan volume overlap.** (A) Glycan N88 in one particular pose. (B) Spatial sampling by glycan N88. Each glycan can take a variety of different conformations and orientations, sampling a large volume in space. (C) Probability distribution map of inter-glycan fractional overlap. Neighboring protomer glycans are indicated by suffix '_2'. (D) Network projected on the Env structure in 2-dimensions. The orange dots indicate the projection of Env protein structure in 2D. Glycan nodes are given by circles, connected by edge lines, colored in blue. (E) The network degree of each node or glycan, given by the number of other nodes it is connected to.

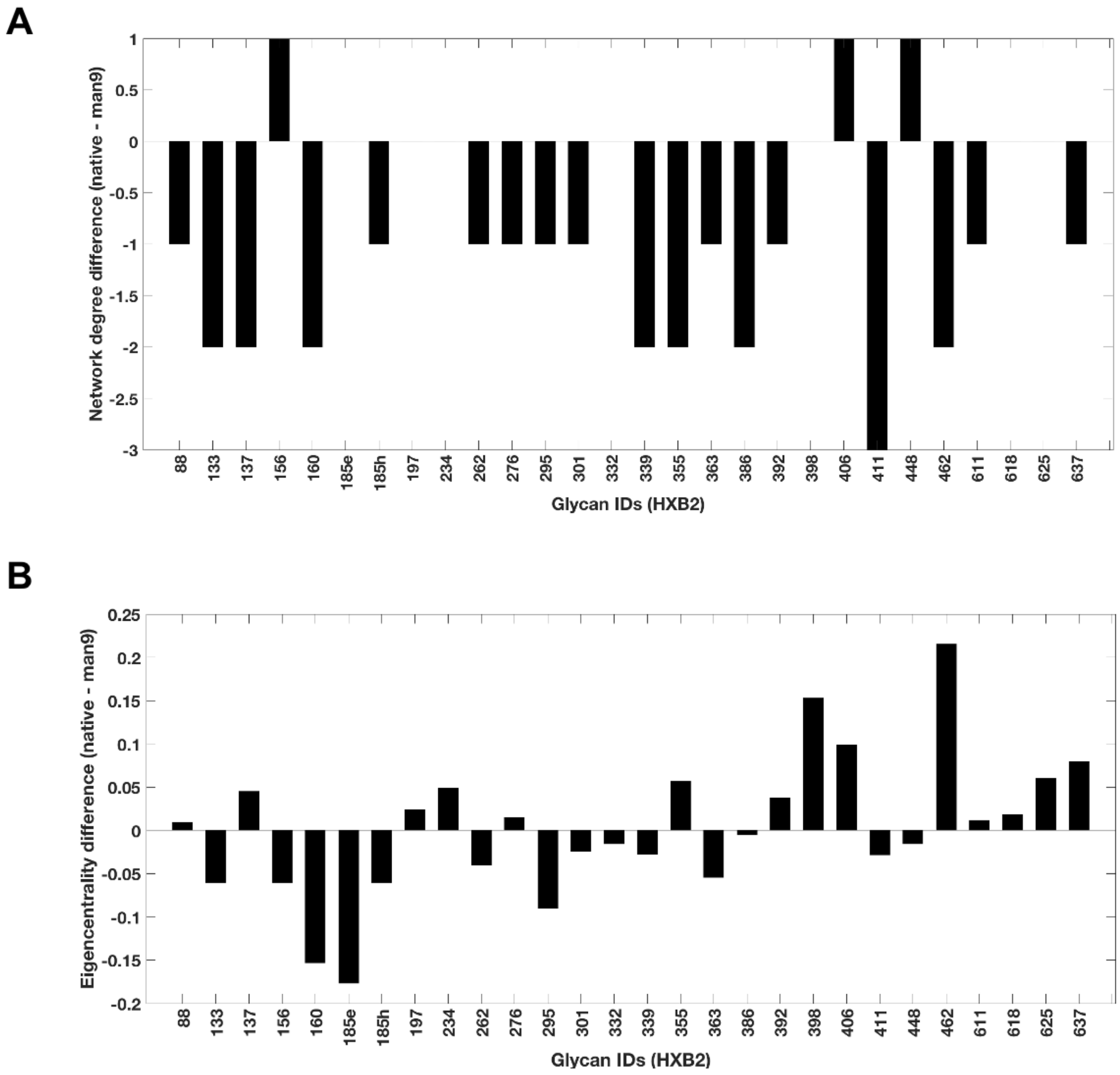

**Figure S4:**

**Network difference between native and all-man9 glycosylation.** Differences are calculated as native minus all-man9. (A) Network degree difference. (B) Eigenvector centrality difference.

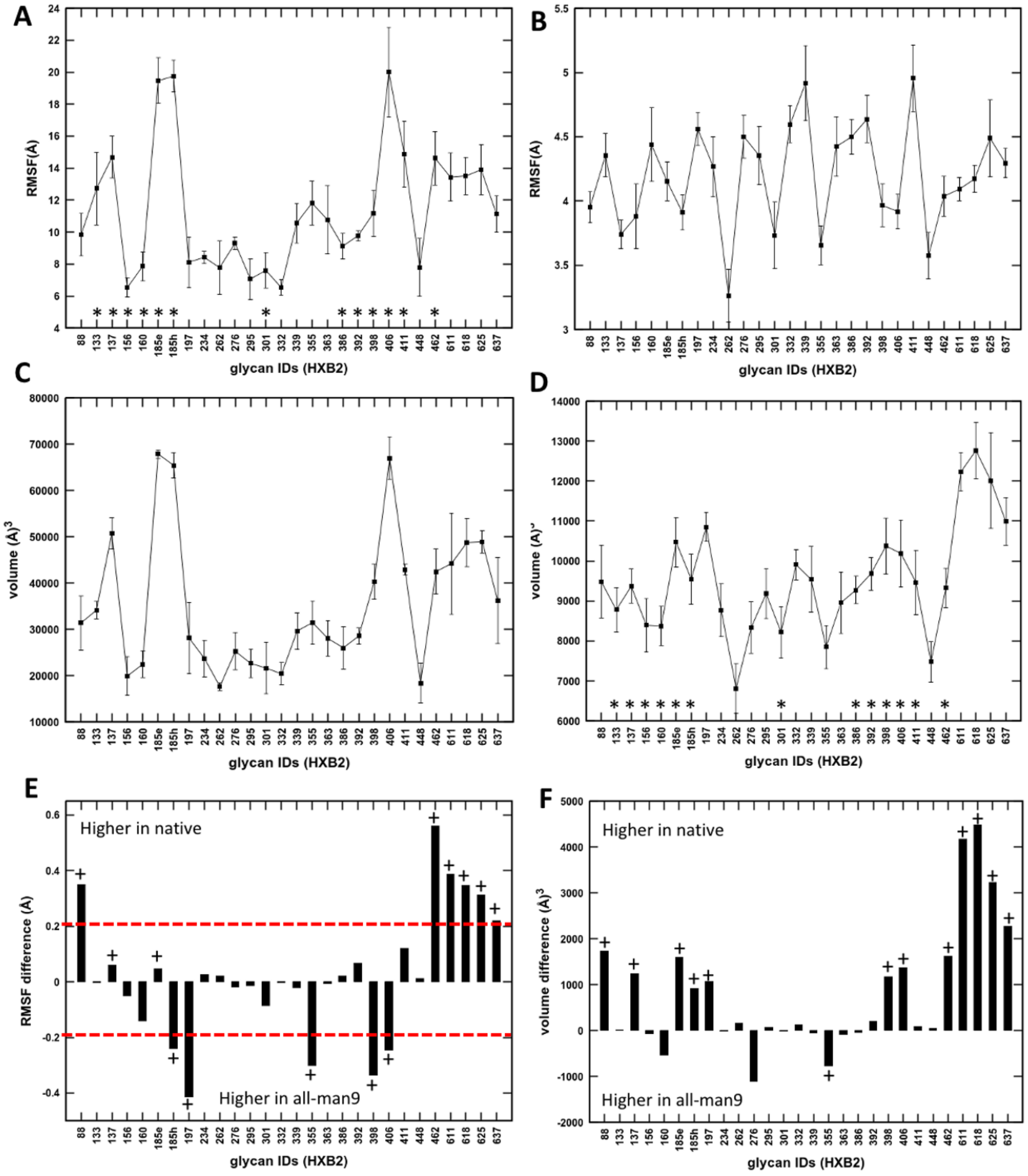

**Figure S5: Structural fluctuations and volume sampling of individual glycans in BG505 native model.** Glycans present in the hyper-variable loop regions of gp120 are indicated by *. Glycans modeled as fucosylated complex or hybrid glycoforms are indicated by +. (A) Site-specific Root Mean Squared fluctuations (RMSF). (B) Reduced RMSF per glycan, where underlying protein backbone are locally aligned to remove the contribution from protein fluctuations. (C) Sampled volume per glycan at each PNGS. (D) Reduced sampled volume per glycan, removing the contribution from protein fluctuations. (E) Difference in RMSF between native and all-man9 model. Native minus all-man9 values are plotted here. (F) Difference in sampled volume between native and all-man9 model. Native minus all-man9 values are plotted here.

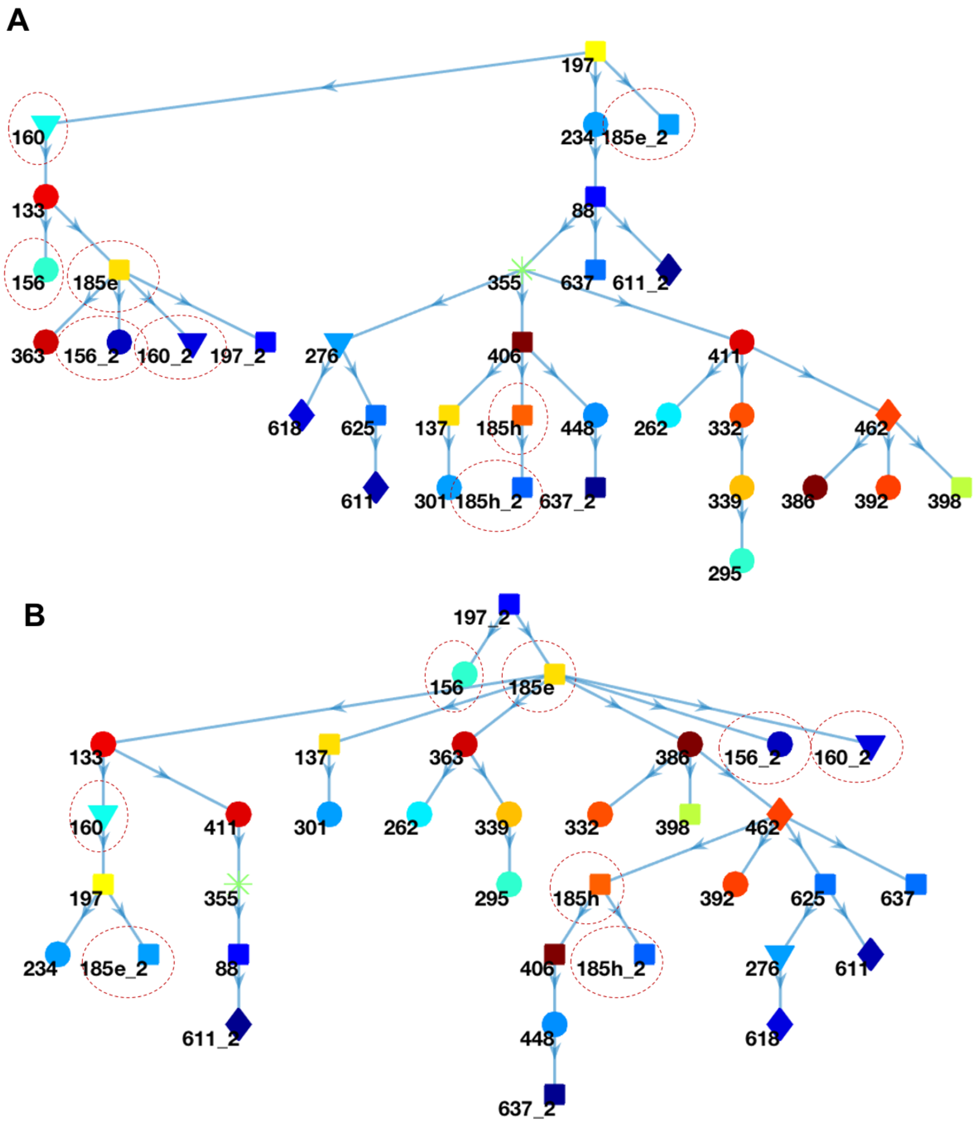

**Figure S6:**

**Glycan 197 directly and indirectly affects glycans in the immediate vicinity of PG9 and PGT145 epitopes.** (A) Shortest paths of communication from glycan N197 to all other glycans in the network. (B) Shortest path of communication from glycan N197 in the neighboring protomer. In both the graphs, those glycans that directly contribute to shielding of PG9 and PGT145 epitopes are circled by red dotted lines.

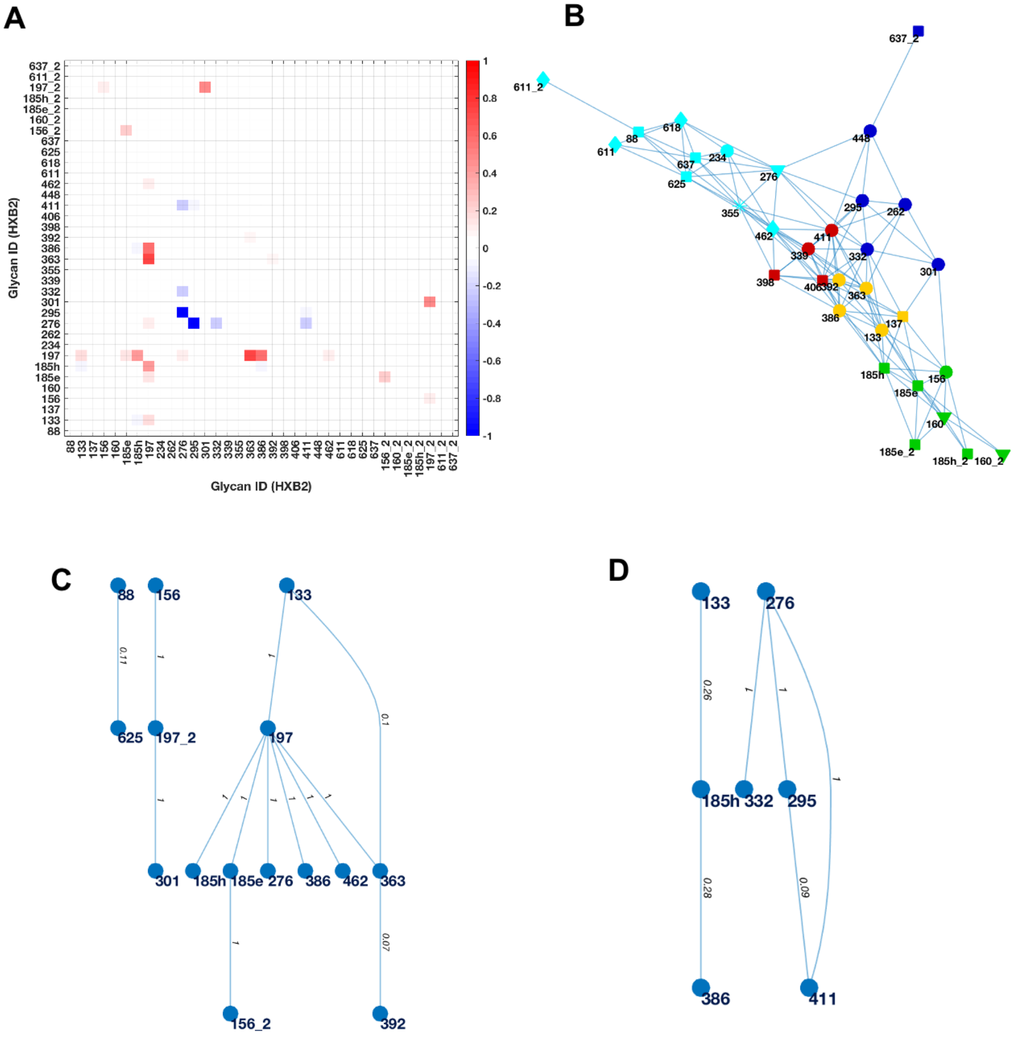

Connectivity loss due to Δ197

Connectivity gain due to Δ197

**Figure S7:**

**Network difference between native glycosylation and deletion of N197 glycan.** (A) Difference in adjacency matrices between the two models, - native minus del197. Red color indicates at least 5% decrease in edge weight, and blue indicates at least 5% increase in edge weight in Δ197 network, as compared to native glycosylation pattern. (B) Network communities in the Δ197 model. (C) Decrease and (D) increase in connectivity due to del197 model in comparison with native glycosylation.

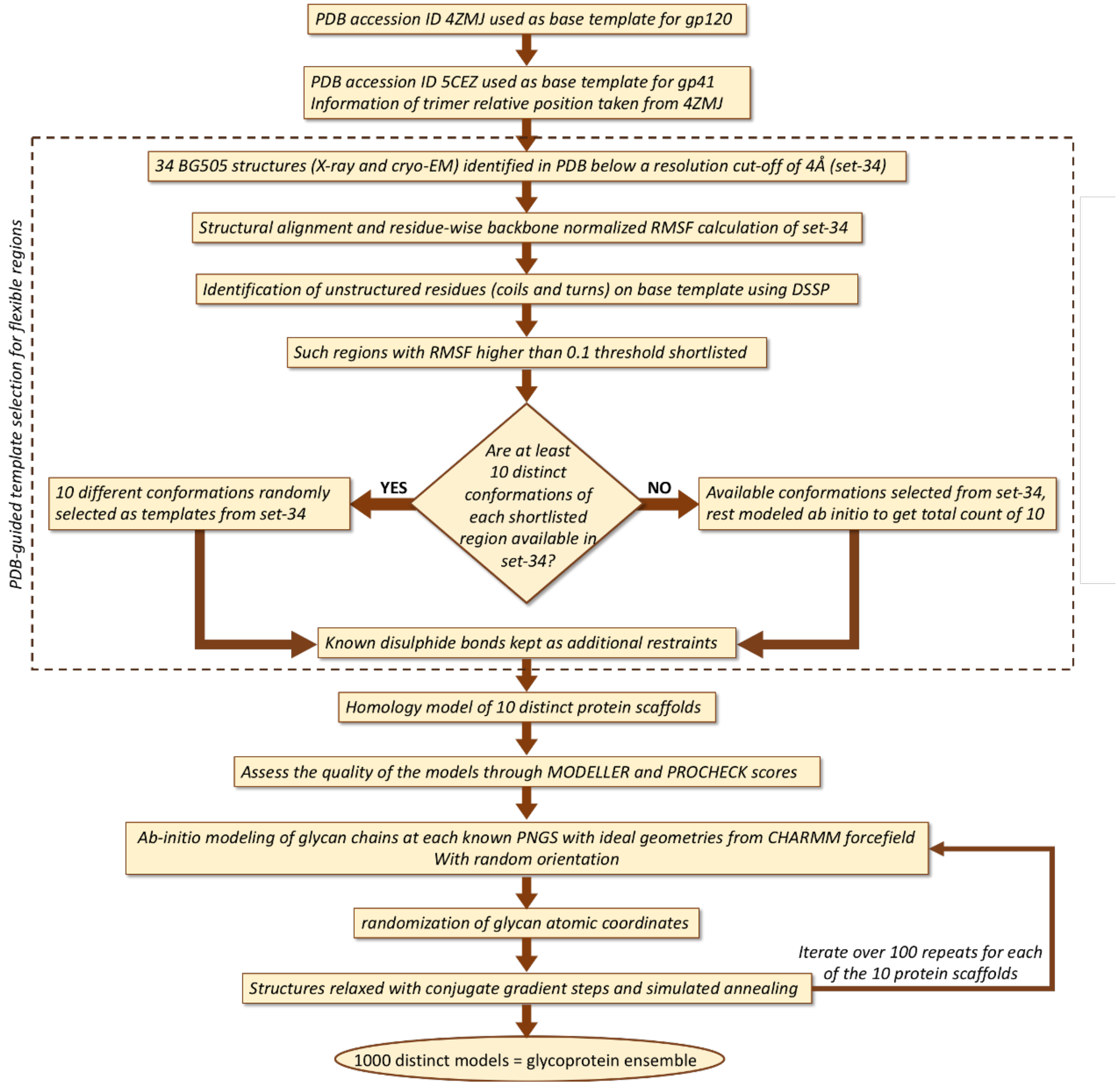

**Figure S8:**

**Workflow describing modeling pipeline for BG505-SOSIP Env glycoprotein ensemble**

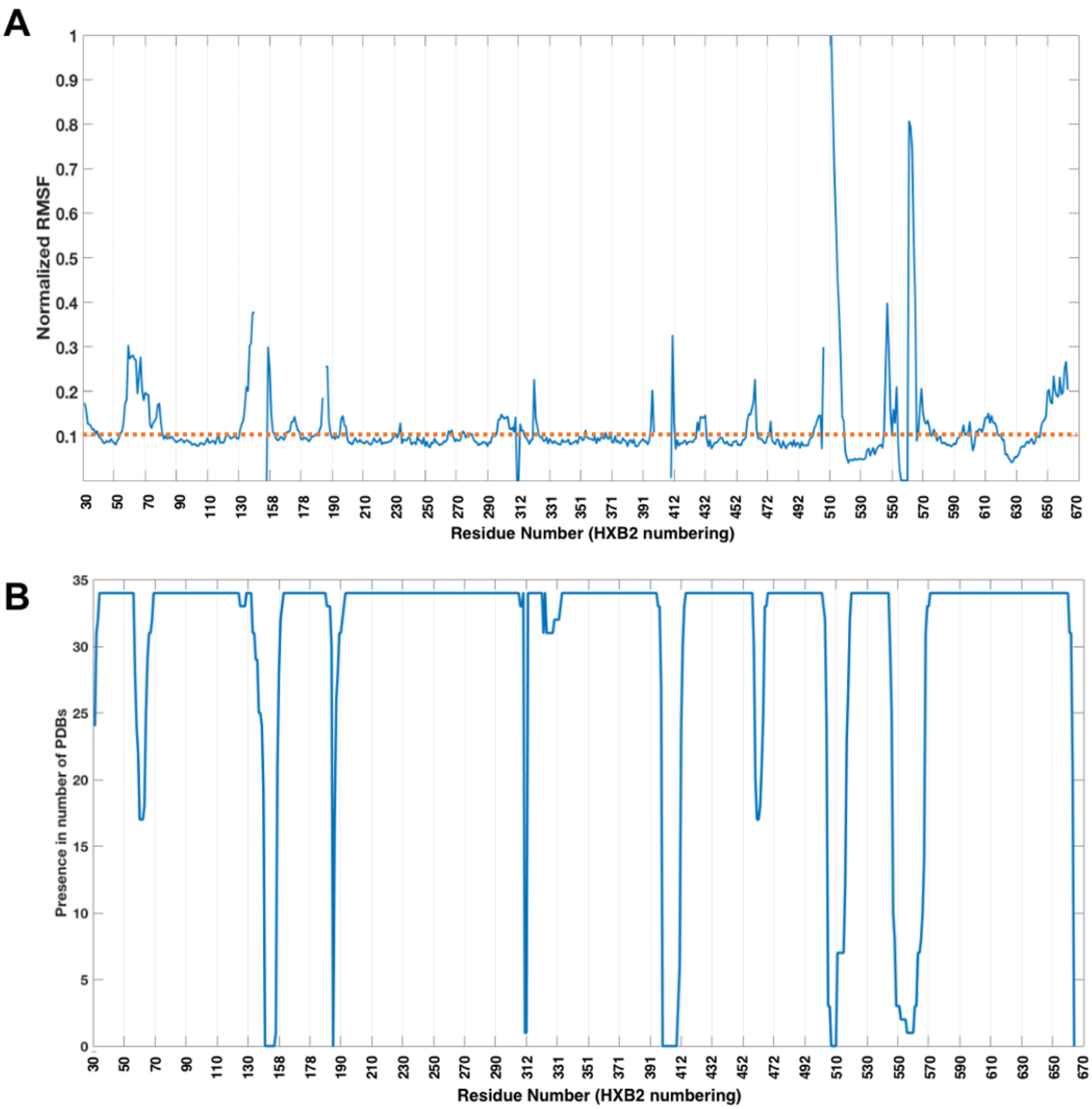

**Figure S9:**

**Homology modeling of unstructured loop regions based on variations in available PDB structures.** (A) Root Mean Squared Fluctuations in protein backbone position based on 34 different PDB structures. Those loop regions having fluctuations higher than the threshold line (dotted, red) were modeled based on 10 randomly selected available datapoints where residues are present. (B) Residue-wise presence in the 34 BG505 PDB candidates. Some of loop regions such as V2 and V4 have missing residues in all available structures.
